# Sleep Spindle-Locked Targeted Memory Reactivation Enhances Declarative Memory Consolidation

**DOI:** 10.64898/2026.05.08.723823

**Authors:** Vaishali Mutreja, Prakriti Gupta, Ovidiu Lungu, Latifa Lazzouni, Ella Gabitov, Habib Benali, Hugo R. Jourde, Giovanni Beltrame, Emily B. J. Coffey, Jean-Marc Lina, Genevieve Albouy, Bradley R. King, Arnaud Boutin, Julie Carrier, Julien Doyon

## Abstract

**Study Objectives:** Sleep spindles are implicated in memory consolidation. Yet direct evidence linking spindle dynamics to declarative memory outcomes remains limited. We thus tested whether targeted memory reactivation (TMR) time-locked to sleep spindles enhances declarative memory, and whether the temporal organization of stimulated spindles–trains versus isolated events–is selectively associated with distinct memory outcomes.

**Methods:** Twenty-eight healthy young adults learned image locations from two categories (animals, clothing) in a grid, each paired with a distinct auditory cue. During overnight NREM sleep, one cue was replayed time-locked to spindles detected in real-time using a closed-loop system (TMR condition); the other served as the non-reactivated control (No-TMR condition). Category-cue assignment was counterbalanced. Post-sleep recall, recognition accuracy, and movement time were assessed.

**Results:** Recall accuracy was significantly higher in the TMR than the No-TMR condition (93.96% vs. 90.61%, *p* = .024), whereas recognition accuracy (*p* = .139) and movement time (*p* = .651) did not differ. Stimulation intensity within spindle trains correlated with the TMR effect on recall (Spearman ρ = .531, *p* = .004), whereas the proportion of isolated spindle stimulations correlated with the TMR effect on recognition (ρ = .563, *p* = .002). Cross-associations were not significant.

**Conclusions:** Spindle-locked TMR enhances recall-based declarative memory retention. The selective association between spindle temporal clustering and memory outcomes suggests that train-embedded and isolated spindles support different aspects of memory consolidation, highlighting spindle temporal context as a functionally relevant dimension of sleep-dependent memory processing.

## Statement of significance

Sleep is essential for stabilizing newly acquired memories. Yet the brain oscillations that support this process during sleep remain poorly understood. Sleep spindles – brief bursts of brain activity that occur during deep sleep stages – have long been linked to memory, though whether their precise timing matters is still unknown. Here we show that re-presenting sounds paired with previously learned declarative material during spindles enhances next-day recall. Moreover, clustered spindles and isolated spindles supported different aspects of memory. These findings clarify how sleep supports memory and open new avenues for spindle-targeted interventions in populations whose memory is compromised by aging or disease.

## Introduction

Sleep plays a critical role in memory consolidation – the process by which newly acquired information is stabilized, integrated with prior knowledge, and transformed into a durable long-term representation ^1–3^. While all sleep stages contribute to this process, the Non-Rapid Eye Movement (NREM) stages, particularly NREM2 and NREM3, have been identified as especially important for declarative memory consolidation, that is, the consolidation of memories for facts and events that can be consciously recalled ^1,4^. During these stages, memory reactivation and integration are supported by the precise temporal coordination of nested electrophysiological events, including hippocampal sharp-wave ripples (80-200 Hz), thalamocortical spindles (11-16 Hz), and slow oscillations (0.5-1 Hz) ^3–8^. Among these, sleep spindles have gained particular attention for their role in enhancing synaptic plasticity and enabling communication between the hippocampus and neocortex, two structures central to memory consolidation ^8,9^.

In support of this view, numerous studies have demonstrated that spindle activity increases following learning and correlates with post-sleep declarative memory performance the next day, underscoring its importance for consolidation ^10–15^. However, identifying the precise spindle features most strongly linked to memory consolidation has remained a challenge, with various properties, including ‘density’ ^13,15–19^, ‘frequency’ ^19–21^, ‘amplitude’ ^22,23^, and ‘power’ ^20,24,25^, each reported as positively associated with post-sleep memory performance. Beyond these conventional features, however, recent work has drawn attention to the temporal organization (i.e., clustering) of spindles, distinguishing spindle trains (sequences of two or more spindles occurring less than 6 s apart) from isolated spindles, which are separated by longer intervals ^8,26^. This distinction is invisible to global sigma power, which integrates activity over time and conflates several spindle properties into a single index ^10,12^. Yet evidence to date suggests that spindle trains contribute selectively to procedural memory consolidation ^8,26–28^, but their role in declarative memory consolidation remains largely unexplored.

The mechanism by which sleep spindles support memory consolidation is thought to involve the reactivation and reprocessing of memory traces encoded during prior wakefulness ^4,5,7,9,29,30^. According to the active systems consolidation framework, spindles play a central mechanistic role by gating the transfer of hippocampal memory representations to neocortical long-term stores through the repeated co-occurrence of sharp-wave ripples and thalamocortical spindles during NREM sleep ^4,7^. Direct evidence for this spindle-linked reactivation mechanism comes primarily from animal studies demonstrating replay of learning-related hippocampal activity during spindle events ^31–37^. Converging evidence has also emerged in humans: Schreiner, Lehmann and Rasch ^38^ showed that memory cues delivered during NREM sleep trigger transient increases in spindle activity, and that disrupting this spindle window immediately after cue delivery abolishes the memory benefit, hence strongly implicating spindle oscillations in the stabilization of reactivated memory traces. Complementing this, Schönauer, Alizadeh, Jamalabadi, Abraham, Pawlizki and Gais ^39^ used multivariate EEG decoding to demonstrate that spindle-band activity (11-16 Hz) during NREM sleep carries material-specific information about prior learning, with the strength of this reprocessing predicting overnight memory consolidation. Collectively, these results establish spindles as a mechanistically relevant carrier of memory reactivation signals in humans, not merely a correlate of the consolidation process. What remains less understood, however, is whether all spindle events contribute equally to reactivation, or whether their temporal organization shapes reactivation efficacy. Spindle trains, by generating multiple successive reactivation windows within a brief interval, may facilitate more iterative hippocampo-cortical communication than isolated spindle events ^8,33^.

Targeted Memory Reactivation (TMR) – a paradigm in which sensory cues paired with to-be-learned material during encoding are re-presented during sleep to trigger reinstatement of cortical representations – provides a direct experimental tool to probe these reactivation processes ^40,41^. In fact, a growing body of evidence indicates that TMR can improve post-sleep memory performance on declarative tasks involving spatial location learning ^42–44^, spatial navigation ^45^, and verbal memory ^46–48^, as reviewed elsewhere ^30,49–53^. Importantly, TMR sensory cues (auditory, olfactory) can be delivered either at predefined times relative to sleep stages (open-loop) or precisely time-locked to specific oscillatory events detected in real time (closed-loop). Most closed-loop TMR work to date has targeted the slow oscillation up-state ^54–56^, with few prior studies examining TMR delivery in relation to sleep spindles ^57,58^. This may be due in part to the long-standing view that auditory processing during spindles was thought to be suppressed ^59,60^. Yet, recent work has revised this view. For example, Jourde and Coffey ^61^ have demonstrated that auditory processing is maintained up to the cortical level during sleep spindles, while Jourde, Sobral, Beltrame and Coffey ^62^, Jourde, Sita, Eyqvelle, Brooks and Coffey ^63^ have further shown that spindle-locked auditory stimulation produces robust neurophysiological effects.

The present study addressed two gaps in the literature. First, we examined whether sleep-dependent declarative memory consolidation is enhanced when TMR cues are delivered time-locked to sleep spindles using a closed-loop approach. To do so, we used a device developed by our group (GB, EC, HJ) called ‘Portiloop’ ^64^, a wearable EEG system with a deep-learning-based spindle detection algorithm, enabling auditory cues to be delivered within milliseconds of online-detected spindle onset. We hypothesized that post-sleep memory performance would be greater for cued (TMR) than uncued (No-TMR) items, consistent with the role of spindles in memory reactivation. Second, building on evidence linking spindle trains to procedural memory consolidation ^8,26^, we explored whether differences in declarative memory consolidation are associated with the temporal clustering of spindles at the time of cue delivery, specifically whether cues coinciding with train-embedded spindles relate differentially to post-sleep memory performance compared to isolated ones. Here, TMR served as the experimental tool to probe spindle temporal organization: by delivering cues time-locked to online-detected spindles and classifying these offline as train-embedded or isolated, we examined whether the temporal context of spindle activity at the moment of cue delivery predicts consolidation outcomes. We hypothesized that memory benefits would be greater when cues coincide with train-embedded spindles, given that trains may constitute a more permissive temporal window for memory reactivation.

## Methods

### 2.1 Participants

Thirty-two healthy young adults (18 females, mean age = 25.01 □ 3.15 years, range = 21–32) were recruited through local advertisement. All participants provided written informed consent prior to participation. The study protocol was approved by the Institutional Review Board at McGill University (22-02-45, A03-B21-22A) and was conducted in accordance with the Declaration of Helsinki (2013). Participants received monetary compensation upon completion of the experiment. All were right-handed, medication-free, non-smokers, and had a body mass index ≤ 25. Participants reported no history of neurological, psychiatric, or sleep disorders and met standard criteria for normal mood, sleep quality, and circadian preference (see Supplementary Section 1.1 for detailed eligibility criteria and screening measures). Participants were excluded if they reported shift work or trans-meridian travel within the three months preceding the study.

Out of 32 participants initially recruited, four were excluded from the final analyses: two due to technical issues during EEG stimulation, one due to photographic memory (disclosed spontaneously during post-session debriefing), and one due to suspected active substance use (identified by experimenter observation during the session). The final sample therefore consisted of 28 participants (see Table 1 for demographic details).

**Table 1.**
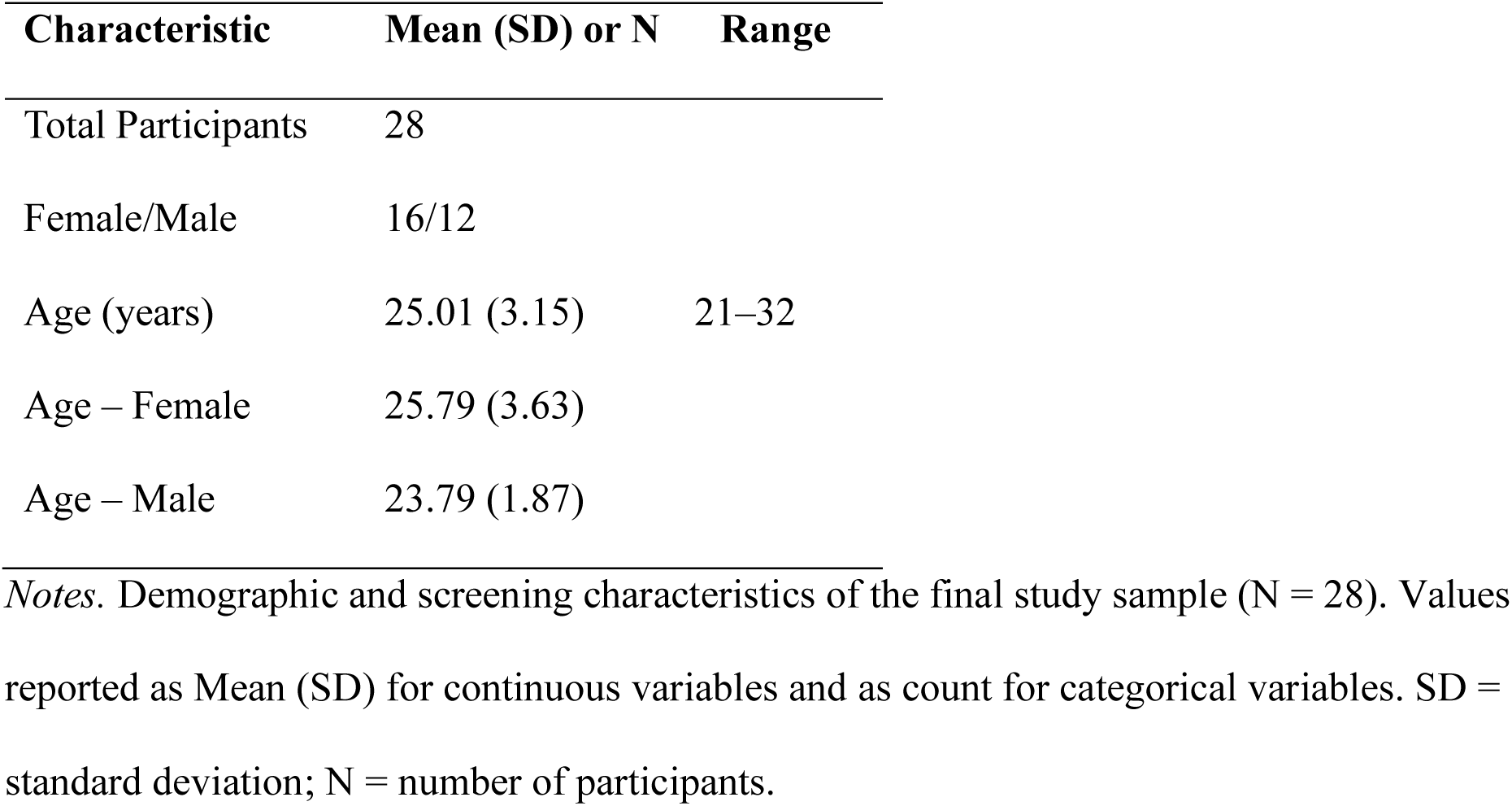
Participants’ characteristics.

### 2.2 General experimental design

The study employed a within-participant design spanning two overnight laboratory visits separated by 4–7 days (Figure 1.a). During the first (habituation) visit, participants acclimated to the sleep laboratory environment and underwent polysomnographic screening by a certified sleep technologist to rule out sleep abnormalities (e.g., sleep apnea, periodic limb movements; Berry et al., 2015). No participants were excluded at this stage; habituation night sleep architecture is summarized in Supplementary Section 2.4 (Table S3).

**Figure 1.**
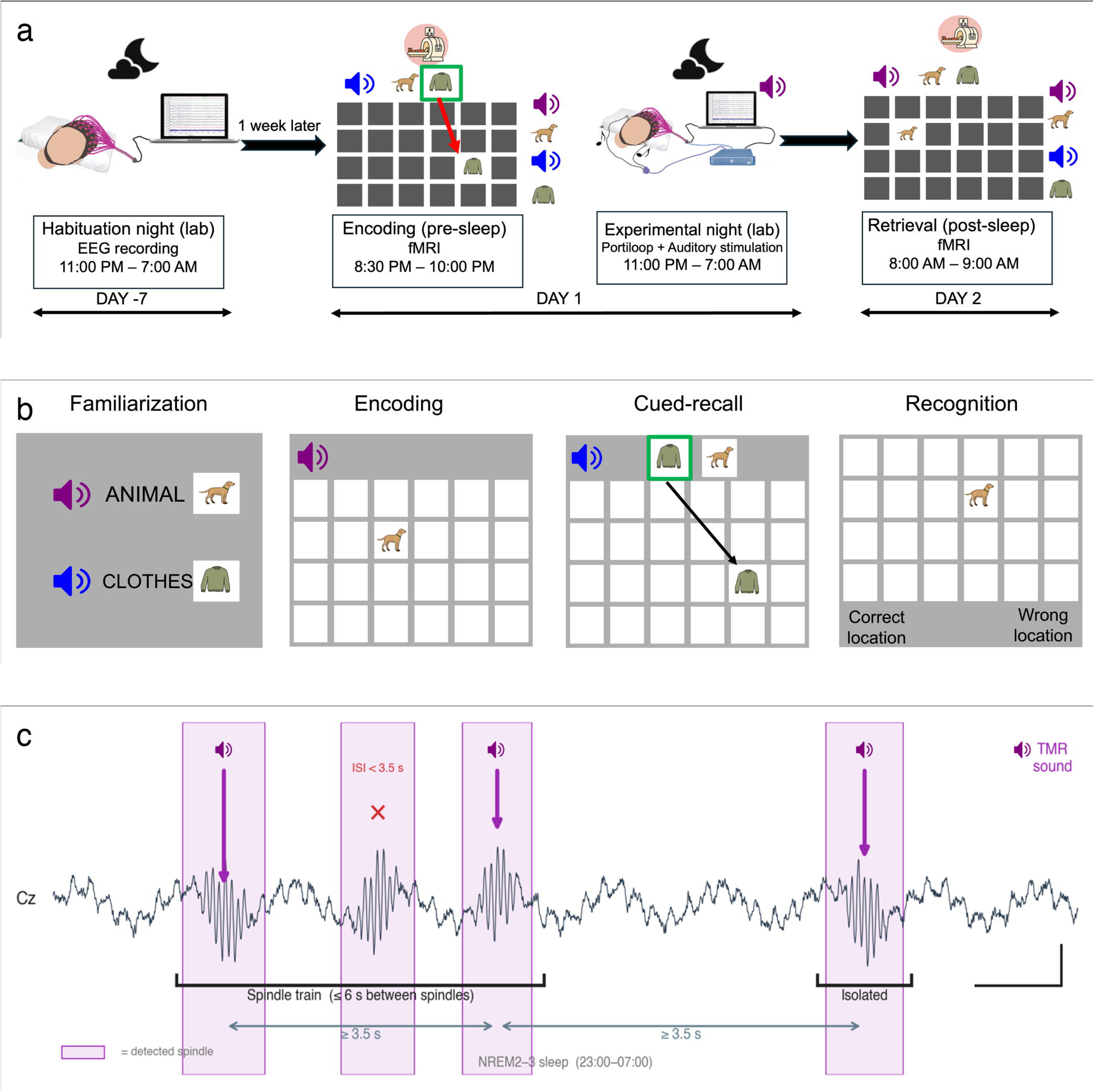
Experimental and task design. (a) Overview of the study protocol. Participants completed a habituation night in the sleep laboratory approximately one week before the experimental visit for polysomnographic screening. Subjects then performed the Object-Spatial Location (OSL) memory task during the evening fMRI session (Day 1), followed by an overnight sleep session with closed-loop spindle-targeted auditory stimulation using the Portiloop device. Memory was assessed again the following morning during a post-sleep fMRI session (Day 2). (b) The Object-Spatial Location (OSL) memory task comprised four phases. During the familiarization phase, participants learned associations between each of two auditory cues and a stimulus category (animals or clothing; counterbalanced across participants). During encoding, participants memorized the spatial locations of 48 images (24 animals, 24 clothing items) arranged in two separate 6 × 4 grids through repeated cued-recall blocks until a learning criterion of ≥15/24 per category was reached. In the cued-recall test, participants heard one of the two sounds, selected the corresponding image from a pair (one per category), and placed it on the grid. Recognition testing (post-sleep only) required participants to distinguish the correct spatial location from an incorrect alternative. (c) Stimulation protocol. During NREM2 and NREM3 sleep (23:00–07:00), the Portiloop device performed real-time spindle detection. Upon detection, the category-associated auditory cue (TMR sound) was delivered via earphones with a minimum inter-stimulus interval of 3.5 s. Items associated with the other sound (No-TMR) received no auditory stimulation. Spindles were classified offline as train-embedded (consecutive spindles ≤ 6 s apart) or isolated (> 6 s from any neighbouring spindle). **Alt text**. Three-panel schematic of the study design. Panel a shows the experimental timeline: a habituation night, an evening fMRI encoding session one week later, an overnight session with closed-loop auditory stimulation, and a post-sleep fMRI retrieval session. Panel b illustrates the four phases of the Object-Spatial Location memory task: familiarization with category-sound pairings, encoding, cued recall, and recognition testing. Panel c shows an example EEG trace from the Cz electrode during NREM 2 to 3 sleep, with three spindles forming a spindle train within 6 seconds of each other and one separate isolated spindle, illustrating the temporal classification of stimulated spindles.

During the experimental visit, participants were required to arrive at the sleep laboratory in the evening (19:30 – 20:00) and were guided to the McConnell Brain Imaging Center (The Neuro, Montreal, Canada). They completed both the encoding and pre-sleep testing phases of the Object-Spatial Location (OSL) declarative memory task while lying supine inside a 3-T Siemens Prisma^Fit^ MRI scanner (see *Task and Stimuli, 2.3*). Following the MRI session, participants returned to the sleep laboratory for overnight EEG monitoring and closed-loop spindle-targeted auditory stimulation using the Portiloop device (Valenchon et al., 2022; see *Polysomnography and TMR Protocol, 2.4*). A post-sleep memory assessment was administered the following morning (07:00 – 09:00) in the MRI scanner.

Full description of the auditory stimuli and calibration procedures, EEG preprocessing as well as spindle detection and classification are provided in the Supplementary Section 1.2–1.3.

### 2.3 Task and stimuli

#### 2.3.1 Object-Spatial Location (OSL) declarative memory task

Declarative memory was assessed using a two-dimensional Object-Spatial Location (OSL) task (Figure 1.b), performed using an MR-compatible trackball response device. The task was adapted and modified from prior TMR paradigms ^41,66,67^ and implemented in PsychoPy ^68^. The task comprised four phases: familiarization, encoding, pre-sleep testing, and post-sleep testing, described in detail below.

Stimuli consisted of 48 colored images selected from the revised Snodgrass and Vanderwart object set ^69^, equally divided into two semantic categories (24 animals, 24 clothing items), selected to be comparable in visual familiarity and complexity across the two categories. The two categories were presented in two separate 6×4 grids – one grid per category – each containing 24 unique spatial locations, such that each image occupied a distinct location within its category-specific grid. Each category was consistently paired with a distinct auditory cue (see 2.3.2 section), which was used for TMR stimulation during sleep. Category-sound assignment was counterbalanced across participants. The auditory cues were selected to avoid any semantic association with the image categories – neither the harmonic tone nor the spoken vowel carries meaning related to animals or clothing.

During *familiarization*, participants were introduced to all 48 images and learned the association between each auditory cue and its corresponding image category. Images were presented category by category: all 24 images from one category were shown first, followed by all 24 images from the other category. The order of categories was counterbalanced across participants. Each image was presented individually for 2000 ms, preceded by the category-specific auditory cue. A category label (e.g., “Animals” or “Clothes”) appeared on screen before each category block, and each image was displayed with its item name printed beneath it (e.g., a picture of a lion with “Lion” below it). Participants were instructed to attend to the images and memorize which auditory cue was paired with which category.

During *encoding*, participants completed repeated blocks consisting of two sequential phases: an image presentation phase and a cued-recall phase. In the *image presentation phase*, images from each category (24 images) were presented one at a time in randomized order within their assigned grid locations (2000 ms per image), each preceded by the corresponding category-specific auditory cue and followed immediately by the other category in the same fashion.

Importantly, both category grids shared the same spatial layout – each grid position appeared once in the animals grid and once in the clothing grid (e.g., the top-left location held a lion in the animals grid and a dress in the clothing grid). Participants therefore learned two distinct spatial maps that overlapped in grid coordinates but were differentiated by category and auditory cue.

In the *cued-recall phase*, each trial began with the presentation of an auditory cue, after which participants were shown two images simultaneously – one randomly selected from each category’s full set of 24 images – and asked to select the one associated with the played cue (“choose image”). Correct selections were indicated by a green border around the image; incorrect selections triggered a red border and the trial advanced. For correctly identified images, participants then indicated the image’s spatial location within the category-specific grid (“choose location”). A correct location response triggered automatic replay of the associated sound, reinforcing the sound-location association for later sleep-based reactivation. At the end of each block, a summary screen displayed the number of correctly recalled locations separately for each category (e.g., “Animals: 15/24 correct – Clothing: 13/24 correct”). Encoding blocks were repeated until participants correctly recalled the locations of at least 15 out of 24 images per category (≥ 62.5%), consistent with encoding criteria established through prior experience with this paradigm in our laboratory, or until a maximum of four blocks was completed, whichever occurred first. Importantly, all participants reached this criterion within four blocks.

Following encoding, participants completed a *pre-sleep cued-recall test* (48 trials, one per image) structured identically to encoding cued-recall phase, but without any feedback. After overnight sleep, memory was reassessed using two tests: a *post-sleep cued-recall test* – also without feedback, comprising 48 trials – and a *recognition test*. In the recognition test, each image was presented twice in silence: once in its correct grid location and once in an incorrect location. Incorrect locations were selected from positions in the same grid that were not assigned to any other image, and the spatial distance between correct and incorrect locations was randomized across trials. Participants indicated for each presentation whether the image was in its correct or incorrect position. The recognition test provided a complementary index of memory distinct from effortful cued recall: whereas recall requires active reconstruction of the spatial location, recognition reflects the ability to discriminate correct from incorrect locations based on familiarity or recollection ^70,71^.

#### 2.3.2 Auditory stimuli

Two auditory stimuli were used throughout the experiment. Both stimuli were 100 ms in duration, sampled at 44.1 kHz, with 10 ms linear onset and offset ramps to prevent audible transients. The stimuli were: (i) a complex harmonic tone with a fundamental frequency of 523 Hz (musical note C5), comprising 12 harmonics with linearly decreasing amplitude, synthetically generated to ensure spectral stability; and (ii) a naturally produced spoken vowel (“A”). The two stimuli were selected to be distinctly recognizable and perceptually dissimilar while remaining comparable in overall loudness following normalization. Neither stimulus carried semantic associations with the image categories used in the task (animals or clothing). Full acoustic construction details are provided in Supplementary Section 1.1.

Each sound was consistently paired with one image category for a given participant, with category-sound assignment counterbalanced across participants.

##### In-scanner calibration

Prior to task onset, individual auditory detection thresholds were established while participants lay supine in the MRI scanner with active scanner sequences running. Tones were initially presented at a baseline volume set to 50% of the maximum output level of the sound delivery system. Participants then reduced the volume until reaching the lowest level at which the cue was clearly audible above scanner background noise, defining their minimum individual threshold. For all subsequent in-scanner phases, the volume was amplified by a consistent fixed factor above this threshold to ensure audibility throughout the session. The amplified level was not systematically recorded as a fixed dB value, as it varied by participant based on their individual threshold; the procedure ensured perceptual adequacy rather than a fixed output level. Sounds were delivered via the S14 Ear Insert system from the Sensimetrics Corporation (https://www.sens.com/), MR-compatible insert earphones during the in-scanner task phases.

##### Sleep-session calibration

For the overnight sleep session, sounds were delivered via Etymotic ER 3C insert earphones with foam tips. Prior to sleep onset, sounds were played to each participant at the candidate stimulation level and participants confirmed that the sounds were audible but would be unlikely to wake them. During overnight stimulation, a trained experimenter continuously monitored the EEG in real time. Whenever an arousal or micro-arousal was observed in close temporal proximity to a stimulus delivery, the volume was reduced in 5 dB steps, and the subsequent stimulation event was monitored to confirm resolution of the arousal response. Volume reductions were required on 8 occasions across the full sample, and no participant required more than two reductions, confirming that stimulation levels were well-tolerated throughout the night. Mean sound pressure levels were 34.62 dB (SD = 24.45) for the harmonic tone and 45.00 dB (SD = 23.15) for the spoken vowel (see Supplementary Section 1.1 for full calibration procedures).

During encoding, sounds preceded image presentation and were replayed following correct spatial responses to reinforce sound-category associations. During sleep, only the auditory cue paired with the TMR image category was delivered during detected spindles, while the other sound was withheld entirely throughout the night (No-TMR condition). Category assignment to TMR vs. No-TMR was counterbalanced across participants.

### 2.4 Polysomnography and TMR protocol

#### 2.4.1 Electroencephalography (EEG) acquisition and online monitoring

Overnight EEG was recorded using the Portiloop v2 system ^64^ at a sampling rate of 250 Hz. The electrode montage comprised four electrodes (Fp1, Fz, Cz, and Pz), placed according to the international 10-20 system, referenced to the left mastoid with the left earlobe as ground.

Offline EEG preprocessing – including broadband filtering (0.3-30 Hz) and artifact rejection using a peak-to-peak amplitude threshold (400 µV; moving window: 200 ms, step: 100 ms) – was implemented in EEGLAB ^72^ using custom MATLAB scripts; full preprocessing details are provided in Supplementary Section 1.2.

During the overnight session, EEG was displayed continuously in 30-second epochs using a 0.5-30 Hz display filter and monitored in real-time – meaning the ongoing EEG signal was observed live on screen by a trained experimenter throughout the night – to identify sleep stages according to standard American Academy of Sleep Medicine criteria ^65^, which defines arousals as abrupt shifts in EEG frequency lasting ≥ 3 seconds. The Cz electrode was used for real-time spindle detection during closed-loop stimulation (see 2.4.2 below). Spindle detection and auditory stimulation were restricted to stable NREM2 and NREM3 sleep and were manually suspended during REM sleep, NREM1, wakefulness, or artifact-related arousals, and restarted only after at least 5 minutes of stable NREM2 sleep.

#### 2.4.2 Closed-loop auditory stimulation and TMR protocol

Closed-loop auditory stimulation was implemented using the Portiloop v2 device ^64^. The Portiloop uses a deep-learning-based artificial neural network (ANN) with convolutional and recurrent units, trained on expert-annotated EEG data from the MODA dataset (Massive Online Data Annotation)^73^ to continuously evaluate the Cz EEG signal and output a spindle confidence score. A spindle detection was triggered when the confidence score exceeded the pre-set probability threshold, at which point an auditory cue was delivered automatically. Given that stimulation was triggered as soon as the classifier confidence score crossed the detection threshold, auditory cues were delivered during the ascending (waxing) phase of the spindle envelope, typically within the first ∼200 ms of spindle onset. A spindle detection confidence threshold of 0.75 was selected based on pilot validation comparing detection performance across multiple threshold values (see Supplementary Section 2.1 for full validation results including precision, sensitivity, and F1-scores; Table S1; Figure S1). This threshold was chosen to favor specificity – ensuring that the majority of cues were delivered during genuine spindle events – while maintaining sufficient sensitivity for adequate stimulation frequency across the night. Overnight Portiloop performance (precision, sensitivity, F1-score) for all 28 participants is reported in Supplementary Section 2.2 (Figure S2).

At the start of the overnight session, one of the two category-specific auditory cues was randomly designated as the TMR cue for that participant’s night, with this assignment counterbalanced across participants. Throughout the overnight stimulation period, whenever the Portiloop detected a spindle, this pre-designated TMR cue was delivered automatically.

Stimulation was applied between 23:00-07:00 and restricted to stable NREM2 and NREM3 sleep, with a minimum inter-stimulus interval (ISI) of 3.5 s between consecutive cue deliveries (Figure 1.c). This ISI was chosen for two reasons. First, it accounts for the post-spindle refractory period, during which new spindle events are suppressed, and memory reactivation is less likely to be effective ^57,74,75^. Second, it was validated empirically through a pilot study comparing memory outcomes for stimulation delivered during spindles versus outside the refractory period (Supplementary Section 2.3, Table S2; Figure S3): stimulation during spindles with a 3.5 s ISI produced more consistent overnight memory improvements than stimulation outside the refractory window, hence supporting the current protocol. Note that because the ISI (3.5 s) was shorter than the train-defining interval (6 s), consecutive spindles within the same train could both receive stimulation, provided they were separated by at least 3.5 s. The image category not associated with the TMR cue received no auditory stimulation during sleep and served as the within-participant No-TMR control condition.

### 2.5 Analyses

#### 2.5.1 Behavioral Analysis

To quantify learning, retention, and sleep-dependent consolidation, behavioral analyses focused on encoding performance, pre- and post-sleep memory outcomes, and other composite measures indexing TMR-related effects.

##### Encoding performance

During *encoding*, performance for each block was quantified as the number of correctly recalled image locations per category (out of 24). Each recall trial involved two sequential steps: participants first identified the image associated with a presented auditory cue (“choose image”), then indicated its spatial location within the category-specific grid (“choose location”). Both steps were required for a trial to be scored as correct. To capture individual differences in learning efficiency, two additional indices were derived: (i) the number of encoding blocks required to reach the learning criterion, and (ii) performance on the final encoding block.

##### Recall accuracy and standardized movement time

Memory performance during pre-and post-sleep cued-recall tests was assessed using two primary variables: (1) *recall accuracy* – the number of correctly recalled image locations per category; and (2) standardized movement time – the time taken to move the cursor to the correct grid location, divided by the Euclidean distance between that location and the initial cursor position at the center of the grid. This normalization controlled for spatial distance effects such that shorter standardized movement times for correct responses were interpreted as reflecting greater spatial memory precision, independent of response speed per se. Standardized movement time was computed for correct trials only and examined both as a TMR vs. No-TMR group-level outcome and as a correlational index in relation to spindle characteristics.

##### Recognition accuracy

To complement recall-based measures with a recognition-based index of memory, a recognition test was administered post-sleep in which each image was presented twice – once in its correct location and once in a randomly selected incorrect location within the same grid (see 2.3.1 section above). Two response types were recorded: (1) *hits* – correct-location trials on which the participant correctly responded, “correct location”; and (2) *correct rejections (CRJ)* – incorrect-location trials on which the participant correctly responded, “wrong location”. Recognition accuracy was defined as: (hits + CRJ)/ 48 x 100, where 48 represents the total number of recognition trials per category (24 correct-location + 24 incorrect-location). This combined measure avoids the ceiling bias that would result from scoring hits alone, as a participant who always responded “correct location” would obtain 100% hits with zero correct rejections.

##### Dependent composite measures

To capture TMR-related effects and individual differences in sleep-dependent consolidation, the following composite measures were used in statistical analyses: (a) *retained items (recall)*: percentage of item locations correctly recalled, computed separately for TMR and No-TMR categories at pre-sleep and post-sleep testing. (b) *Recognition accuracy*: percentage of recognition trials answered correctly – (hits + CRJ)/ 48 x 100 – computed separately for TMR and No-TMR categories at post-sleep testing. (c) *TMR effect*: the difference between TMR and No-TMR conditions, computed for (i) *recall*: difference in the percentage of retained items; (ii) *recognition*: difference in the percentage of recognition accuracy scores. (d) *Change in standardized movement time*: overnight change (post-sleep minus pre-sleep) in standardized movement time for correct responses, compared between TMR and No-TMR conditions.

##### Statistical analysis

Behavioral statistical analyses were conducted using linear mixed-effects models (LMEs) and paired-sample comparisons implemented in SPSS (Statistical Package for Social Sciences, version 29.0.0.0) and MATLAB (R2023b: MathWorks). LMEs were used for encoding analyses to account for the repeated-measures structure across blocks while accommodating individual variability in learning trajectories; all LME models used restricted maximum likelihood estimation and Satterthwaite approximation for degrees of freedom, with random intercepts for participants, and estimated marginal means were compared using Bonferroni-corrected pairwise tests.

To test the primary hypothesis that spindle-targeted TMR enhances memory consolidation, paired-sample *t*-tests compared TMR and No-TMR conditions for retained item locations, recognition accuracy, and overnight change in movement time. Normality of the difference scores was evaluated using Shapiro-Wilk tests; all three comparisons met the normality assumption (*p*s > .05). Wilcoxon signed-rank tests are additionally reported as non-parametric robustness checks. Because recall accuracy, recognition accuracy, and movement time index distinct aspects of memory performance, these planned comparisons were evaluated at α = .05 without correction for multiple comparisons. Effect sizes (Cohen’s *d* for paired *t-*tests, Spearman’s ρ for correlations) are reported alongside *p*-values to facilitate interpretation.

#### 2.5.2 EEG data Analysis

Overnight EEG data from all 28 participants were analyzed using MATLAB (R2023b; MathWorks) and EEGLAB ^72^. Sleep stages were scored manually according to standard AASM criteria ^65^ by a registered polysomnographic technologist using the *Counting Sheep PSG* toolbox^77^. EEG preprocessing – including broadband filtering (0.3-30 Hz) and artifact rejection using a peak-to-peak amplitude threshold (400 µV; moving window: 200 ms, step: 100 ms) – was applied to all 28 recordings; full preprocessing details are provided in Supplementary Section 1.2.

##### Spindle detection parameter optimization

To tune the parameters of the two automated spindle detection algorithms, the same registered sleep expert who scored sleep stages for all 28 participants also manually annotated sleep spindles in EEG recordings from three of those participants, selected to represent a broad range of sleep architectures and spindle morphologies. These expert annotations served as reference standard for parameter optimization only and were distinct from the final ground-truth spindle set used in analyses (described below).

Parameters were optimized independently for each algorithm by comparing algorithmic detections against the expert annotations and balancing sensitivity, precision, and F1 score to achieve the best trade-off for robust spindle detection. For the Lacourse A7 detector, four parameters were adjusted: absolute sigma power, relative sigma power, covariance, and correlation between sigma-band and broadband signals, following the parameter selection framework described in Lacourse, Yetton, Mednick and Warby ^73^. For the Wamsley detector, a single parameter was tuned: the relative amplitude threshold of sigma power, adjusted to minimize false positives. The optimized parameter set for each algorithm was subsequently applied to all 28 participants’ recordings.

##### Offline spindle detection pipeline

Following preprocessing, the tuned algorithms were applied to all 28 participants’ EEG recordings to detect spindles across NREM2 and NREM3 sleep (Figure 2). Spindle detection was restricted to periods of stable NREM2 and NREM3 sleep as identified during real-time sleep staging. Both the Wamsley and Lacourse A7 algorithms were run independently, after which overlapping detections – within each algorithm and across the two algorithms – were identified and merged into a single candidate spindle set.

**Figure 2.**
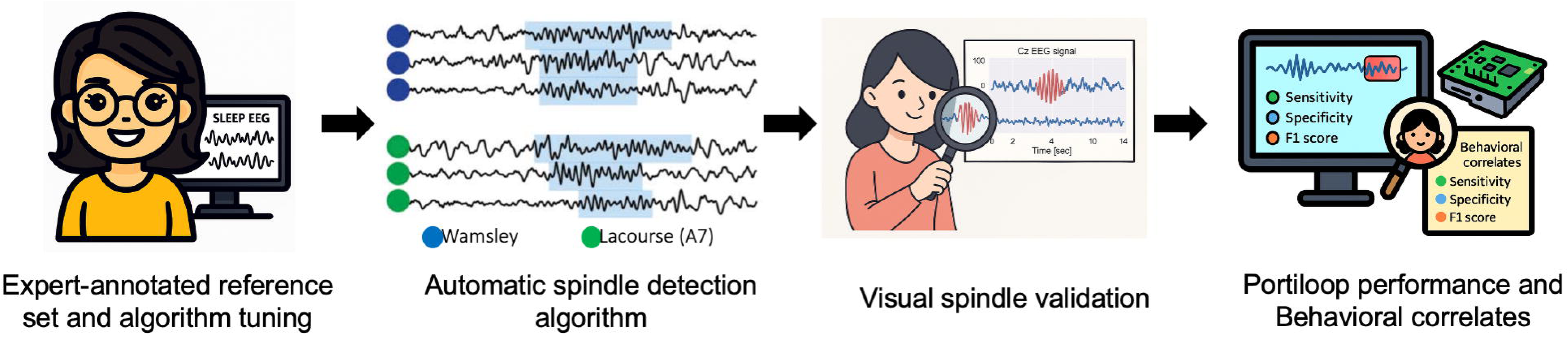
Offline spindle detection and validation pipeline. Expert-annotated sleep EEG from a representative subset of three participants (selected to capture a broad range of sleep architectures and spindle morphologies) was used to optimize spindle detection parameters for each algorithm independently. Two complementary detection algorithms (Lacourse et al., 2020; Wamsley et al., 2012) were applied to EEG recordings from all 28 participants to identify candidate spindles across NREM2 and NREM3 sleep. Detected events were merged across algorithms and subjected to visual inspection to confirm detection accuracy and exclude false positives. Only visually validated spindles were retained as ground-truth (GT) spindles for subsequent analyses. **Alt text**. Four-panel illustrated pipeline for offline spindle detection and validation. Panel one shows expert annotation of a reference EEG dataset for algorithm tuning. Panel two shows automatic spindle detection by two algorithms (Wamsley and Lacourse) producing candidate spindle markers on multi-channel EEG traces. Panel three shows visual validation by a trained scorer using a magnifying glass on a Cz EEG signal. Panel four shows performance metrics including sensitivity, specificity, and F1 score evaluated against the validated ground truth.

All candidate spindle events then underwent visual inspection by a trained researcher to confirm detection accuracy. Spindles were rejected if they: (i) occurred within EEG segments contaminated by artifact, (ii) lacked the characteristic waxing-and-waning amplitude modulation of genuine sleep spindles, (iii) exhibited atypical duration, or (iv) reflected broadband noise rather than spindle-specific sigma activity. Candidate and GT spindle counts are reported in Table 2. Only GT spindles were retained for all subsequent analyses.

**Table 2.**
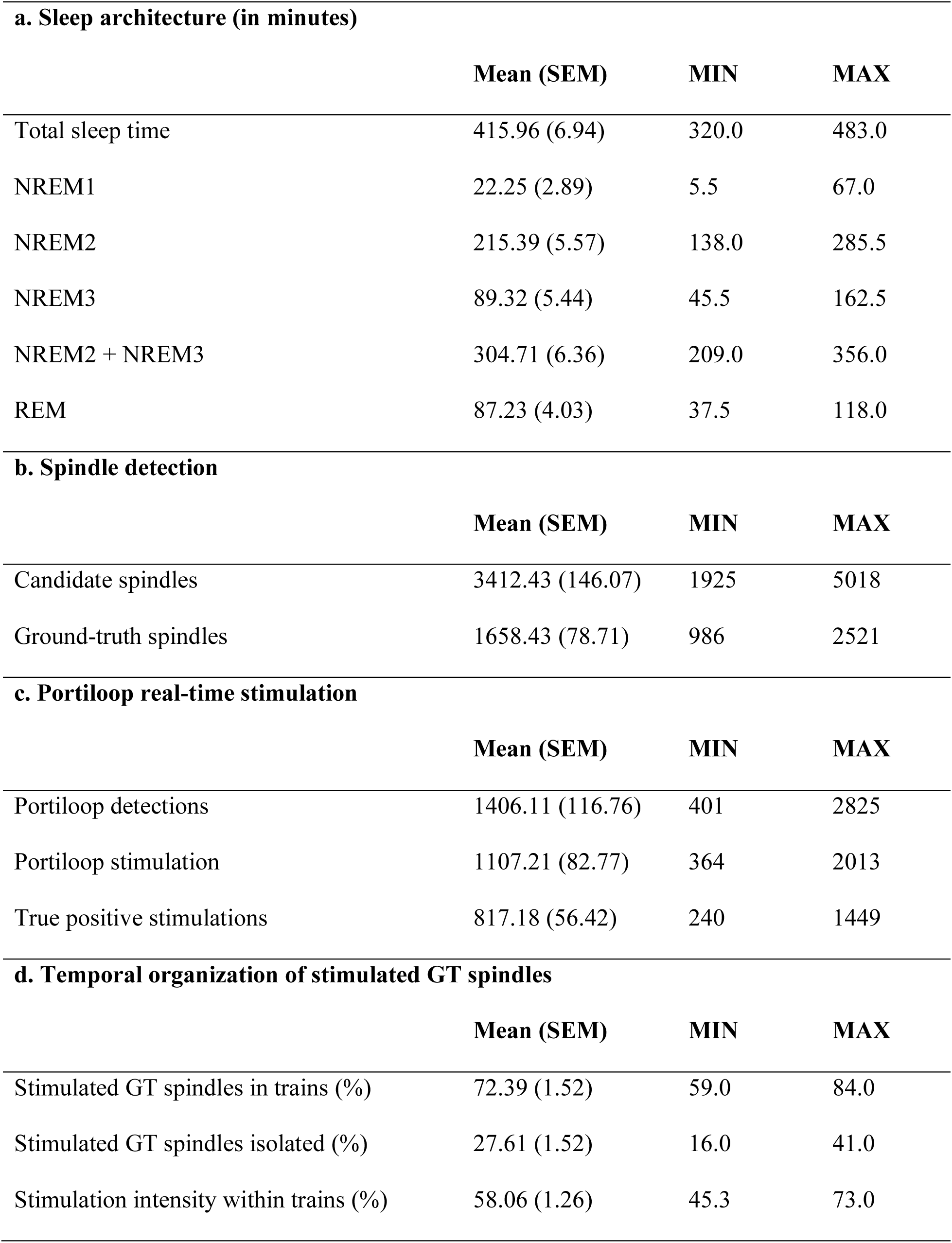
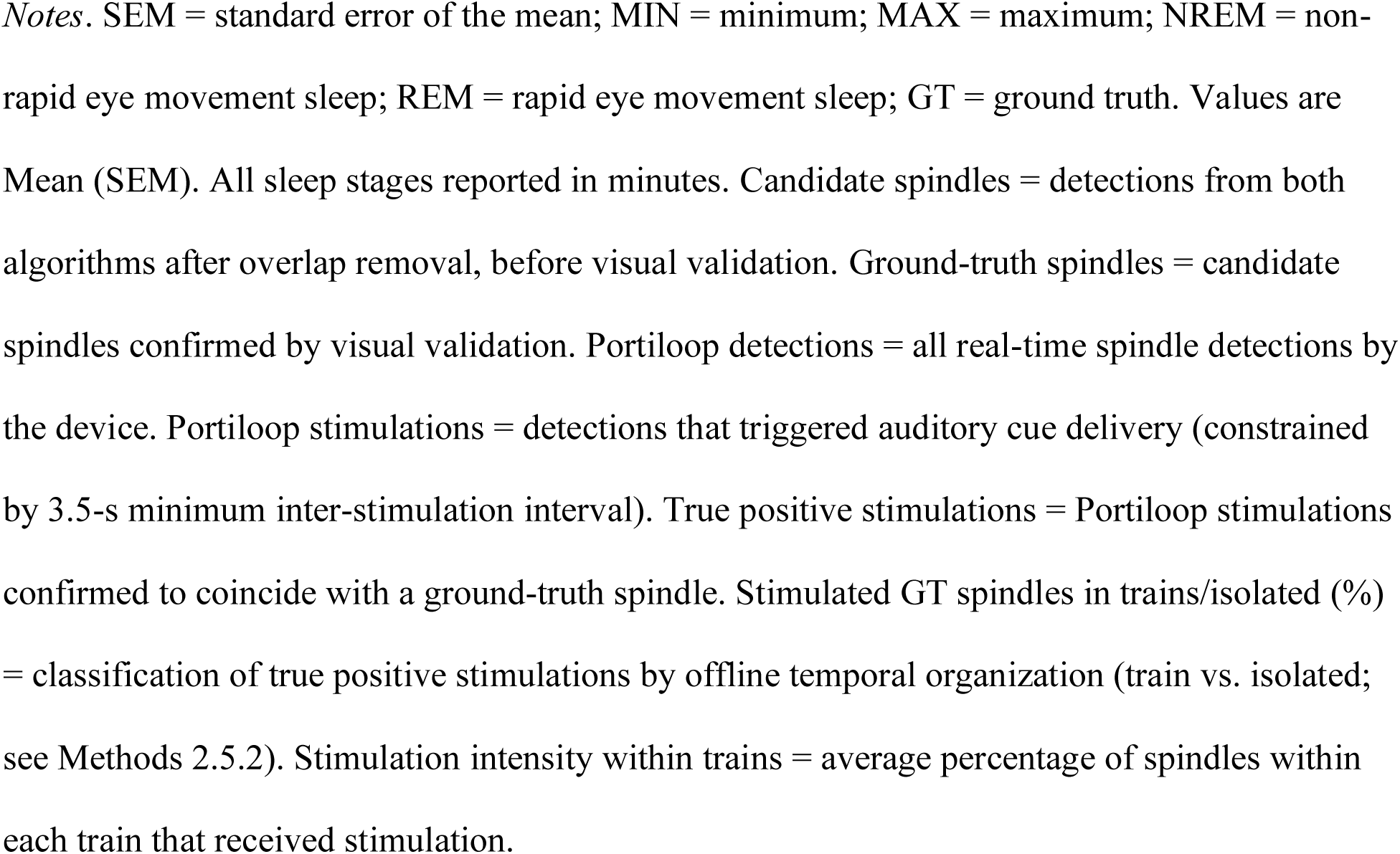
Sleep architecture and sleep spindle characteristics.

| **a. Sleep architecture (in minutes)** |  |  |  |
| --- | --- | --- | --- |
|  | **Mean (SEM)** | **MIN** | **MAX** |
| Total sleep time | 415.96 (6.94) | 320.0 | 483.0 |
| NREM1 | 22.25 (2.89) | 5.5 | 67.0 |
| NREM2 | 215.39 (5.57) | 138.0 | 285.5 |
| NREM3 | 89.32 (5.44) | 45.5 | 162.5 |
| NREM2 + NREM3 | 304.71 (6.36) | 209.0 | 356.0 |
| REM | 87.23 (4.03) | 37.5 | 118.0 |
| **b. Spindle detection** |  |  |  |
|  | **Mean (SEM)** | **MIN** | **MAX** |
| Candidate spindles | 3412.43 (146.07) | 1925 | 5018 |
| Ground-truth spindles | 1658.43 (78.71) | 986 | 2521 |
| **c. Portiloop real-time stimulation** |  |  |  |
|  | **Mean (SEM)** | **MIN** | **MAX** |
| Portiloop detections | 1406.11 (116.76) | 401 | 2825 |
| Portiloop stimulation | 1107.21 (82.77) | 364 | 2013 |
| True positive stimulations | 817.18 (56.42) | 240 | 1449 |
| **d. Temporal organization of stimulated GT spindles** |  |  |  |
|  | **Mean (SEM)** | **MIN** | **MAX** |
| Stimulated GT spindles in trains (%) | 72.39 (1.52) | 59.0 | 84.0 |
| Stimulated GT spindles isolated (%) | 27.61 (1.52) | 16.0 | 41.0 |
| Stimulation intensity within trains (%) | 58.06 (1.26) | 45.3 | 73.0 |

*Notes.* SEM = standard error of the mean; MIN = minimum; MAX = maximum; NREM = non-rapid eye movement sleep; REM = rapid eye movement sleep; GT = ground truth. Values are Mean (SEM). All sleep stages reported in minutes. Candidate spindles = detections from both algorithms after overlap removal, before visual validation. Ground-truth spindles = candidate spindles confirmed by visual validation. Portiloop detections = all real-time spindle detections by the device. Portiloop stimulations = detections that triggered auditory cue delivery (constrained by 3.5-s minimum inter-stimulation interval). True positive stimulations = Portiloop stimulations confirmed to coincide with a ground-truth spindle. Stimulated GT spindles in trains/isolated (%) = classification of true positive stimulations by offline temporal organization (train vs. isolated; see Methods 2.5.2). Stimulation intensity within trains = average percentage of spindles within each train that received stimulation.

##### Spindle clustering

GT spindles were classified, based on their temporal organization, into *trains* and *isolated* spindles following criteria described by Boutin and Doyon ^8^. A spindle train was defined as a sequence of two or more consecutive spindles each separated by no more than 6 s from the preceding or subsequent spindle. Isolated spindles were those separated by more than 6 s from both the preceding and the subsequent spindle. The 6-s criterion is grounded in empirical evidence that infraslow (∼0.02 Hz) oscillatory dynamics organize spindle clustering on a timescale of approximately 3-6 seconds ^78,79^, with spindles separated by more than 6 s showing distinct temporal and functional properties relative to those occurring within this window ^26^. Auditory stimulation events were then mapped onto this ground-truth classification: each stimulated spindle was labelled as occurring within a train or in isolation according to its offline-defined spindle context. Critically, a stimulated spindle was classified as train-embedded even if adjacent spindles within the same train were not stimulated – because train membership reflects intrinsic spindle dynamics, not stimulation timing.

##### Variables included in EEG analyses

The following spindle-related variables were extracted for each participant: (1) *GT spindles*: total number of visually validated spindles from offline detection across NREM2 and NREM3 sleep. (2) *Stimulated spindles*: total number of real-time Portiloop stimulations, including both true positives and false positives. (3) *True positive stimulations*: the subset of stimulated spindles confirmed as GT spindles – equivalent to the true positive count from the overlap between real-time detections and offline GT annotations. (4) *Train-stimulated GT spindles*: true positive stimulations occurring within offline-defined spindle trains. (5) *Isolated stimulated GT spindles*: true positive stimulations occurring as offline-defined isolated spindle events.

Two derived proportion measures were also computed to examine the relationship between spindle temporal organization and memory performance: (a) *Proportion of GT stimulations within trains*: the number of train-stimulated GT spindles divided by the total number of true positive stimulations, reflecting the relative contribution of train-embedded stimulation events to the overall GT stimulation set. (b) *Proportion of GT stimulations in isolation*: the number of isolated stimulated GT spindles divided by the total number of true positive stimulations. By construction, measures (a) and (b) are complementary proportions summing to 1. (*c*) *Stimulation intensity within trains*: for each spindle train, the number of stimulated GT spindles within that train divided by the total number of GT spindles in that train; values were then averaged across all trains for each participant. This measure indexes how densely each train was sampled by the closed-loop stimulation protocol. Unlike measures (a) and (b), which classify stimulated spindles, measure (c) captures the within-train stimulation rate and is the primary predictor variable in the correlational analyses. All spindles related measures were computed with respect to visually validated GT spindles only.

### Statistical analyses

Descriptive statistics were computed for sleep architecture variables and spindle characteristics during NREM2 and NREM3 sleep at Cz electrode. To examine whether reactivation processes differed between spindle types, Pearson correlations were computed between each proportion measure (proportion of GT stimulations within trains; proportion of GT stimulations in isolation) and each TMR-related behavioral outcome (TMR effect on recall, TMR effect on recognition accuracy, and TMR effect on standardized movement time), yielding four primary correlations. Because the proportion of isolated stimulations showed evidence of non-normality (Shapiro-Wilk *W* = .919, *p* = .033), Spearman rank correlations were computed as the primary analysis, with Pearson correlations reported alongside for completeness. Both methods yielded equivalent conclusions. 95% confidence intervals for correlation coefficients were computed using Fisher’s z transformation. To test whether the association between spindle type and memory outcome differed significantly between train-embedded and isolated spindles, Fisher r-to-z transformations were applied to compare corresponding Pearson correlation coefficients across spindle types. All statistical analyses were conducted in MATLAB (R2023b; MathWorks) and SPSS (version 29.0.0.0). Statistical significance was set at *p* < .05.

## Results

### Behavioral results

#### Encoding: Block-wise memory performance

To assess encoding of item locations across the two stimulus categories, a linear mixed-effects model was used with Block and Condition (items later assigned to the TMR vs. No-TMR condition) as within-subject factors. Because encoding followed a criterion-based procedure (≥ 15/24 correct per category), the number of encoding blocks varied across participants (range: 1-4), and a linear mixed-effects model was used to accommodate the resulting unbalanced design.

This analysis revealed a significant main effect of Block, *F*(3, 106.08) = 133.79, *p* < .001, indicating that recall accuracy improved across successive encoding blocks. Crucially, there was no main effect of Condition, *F*(1, 37.04) = 0.26, *p* = .61, indicating that encoding performance did not differ between item categories later assigned to the TMR and No-TMR conditions. The Block x Condition interaction was also not significant *F*(3, 104.25) = 0.49, *p* = .69, suggesting that learning trajectories were comparable for item categories later assigned to the two conditions (Figure 3). Note that stimulus category (animals, clothing) was counterbalanced across the TMR and No-TMR conditions; the present analysis therefore tested directly whether items later reactivated during sleep differed from non-reactivated items at baseline.

**Figure 3.**
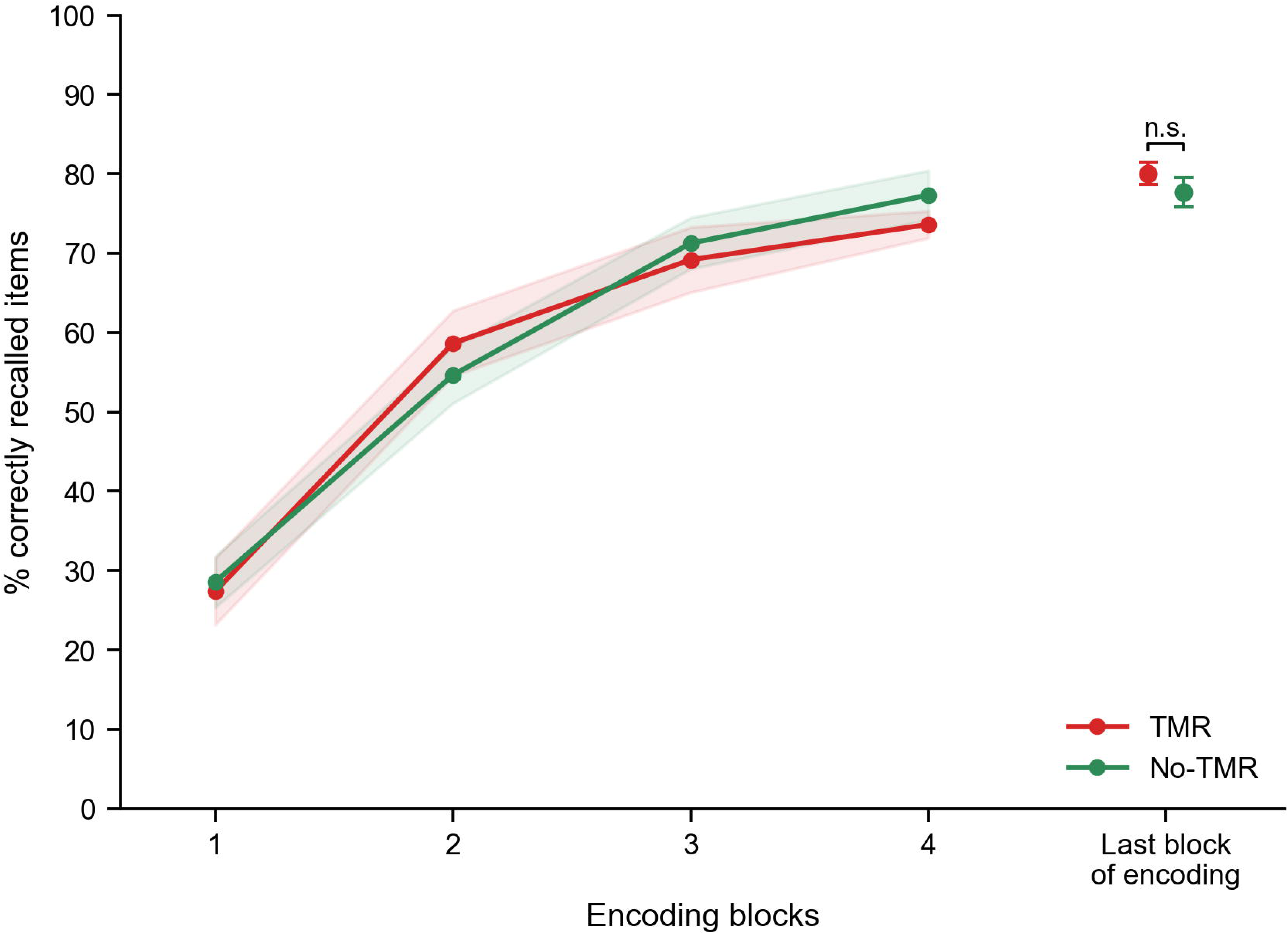
Encoding performance. Percentage of item locations correctly recalled across encoding blocks for item categories later assigned to the TMR (red) and No-TMR (green) conditions. Lines represent group means and shaded areas represent ±1 SEM. The bracket at the final encoding block indicates the paired-samples *t*-test comparing conditions (n.s., *p* = .23). **Alt text**. Line plot of encoding performance across four blocks. The y-axis shows percentage of item locations correctly recalled, ranging from approximately 30 percent at block 1 to approximately 75 percent at block 4. Two overlapping curves show the TMR and No-TMR conditions following nearly identical trajectories with shaded standard error bands. A separate pair of points with error bars on the right indicates the comparison at the final encoding block, marked non-significant.

To further confirm equivalent encoding strength at the end of the encoding phase, a paired samples *t*-test was conducted on performance during each participant’s final encoding block, comparing items categories later assigned to the TMR (*M* = 19.21, *SEM* = .33) and No-TMR (*M* = 18.64, *SEM* = .43) conditions. No significant difference was found, *t*(27) = 1.22, *p* = .23, confirming comparable encoding performance across conditions. An additional analysis confirmed that encoding performance did not differ between stimulus categories (animals vs. clothing; Supplementary Section 2.5, Table S4)

Together, these results indicate that encoding strength and learning dynamics did not differ between conditions prior to sleep. This equivalence minimizes the possibility that any post-sleep differences observed between the TMR and No-TMR conditions would reflect differences in encoding difficulty or baseline performance rather than sleep-dependent consolidation processes.

### TMR time-locked to sleep spindles enhance memory performance

Paired samples *t*-tests were conducted to assess whether post-sleep memory performance differed between the TMR and No-TMR conditions. Recall performance, measured as the percentage of item locations retained from pre-to post-sleep, was significantly higher in the TMR condition (*M* = 93.96%, *SEM* = 0.94) than the No-TMR condition (*M* = 90.61%, *SEM* = 1.17), *t*(27) = 2.39, *p* = .024, *d* = 0.45. A Wilcoxon signed-rank test confirmed this result (*W* = 90, *p* = .030), corroborating the finding under a non-parametric framework.

Recognition accuracy, measured as the percentage of correct responses across hits and correct rejections [(hits + correct rejections)/48], did not differ significantly between the TMR (*M* = 92.26%, *SEM* = 1.08) and No-TMR conditions (*M* = 90.48%, *SEM* = 1.37), *t*(27) = 1.52, *p* = .139, *d* = 0.29. Overnight change in movement time for correctly recalled item locations also did not differ between the TMR (*M* = 86.54 ms, *SEM* = 34.77) and No-TMR conditions (*M* = 68.51 ms, *SEM* = 29.69), *t*(27) = 0.46, *p* = .651, *d* = 0.09. Together, these results (Figure 4) provide evidence that TMR time-locked to sleep spindles selectively enhanced recall-based memory consolidation, whereas recognition accuracy and response speed were not significantly modulated by the intervention.

**Figure 4.**
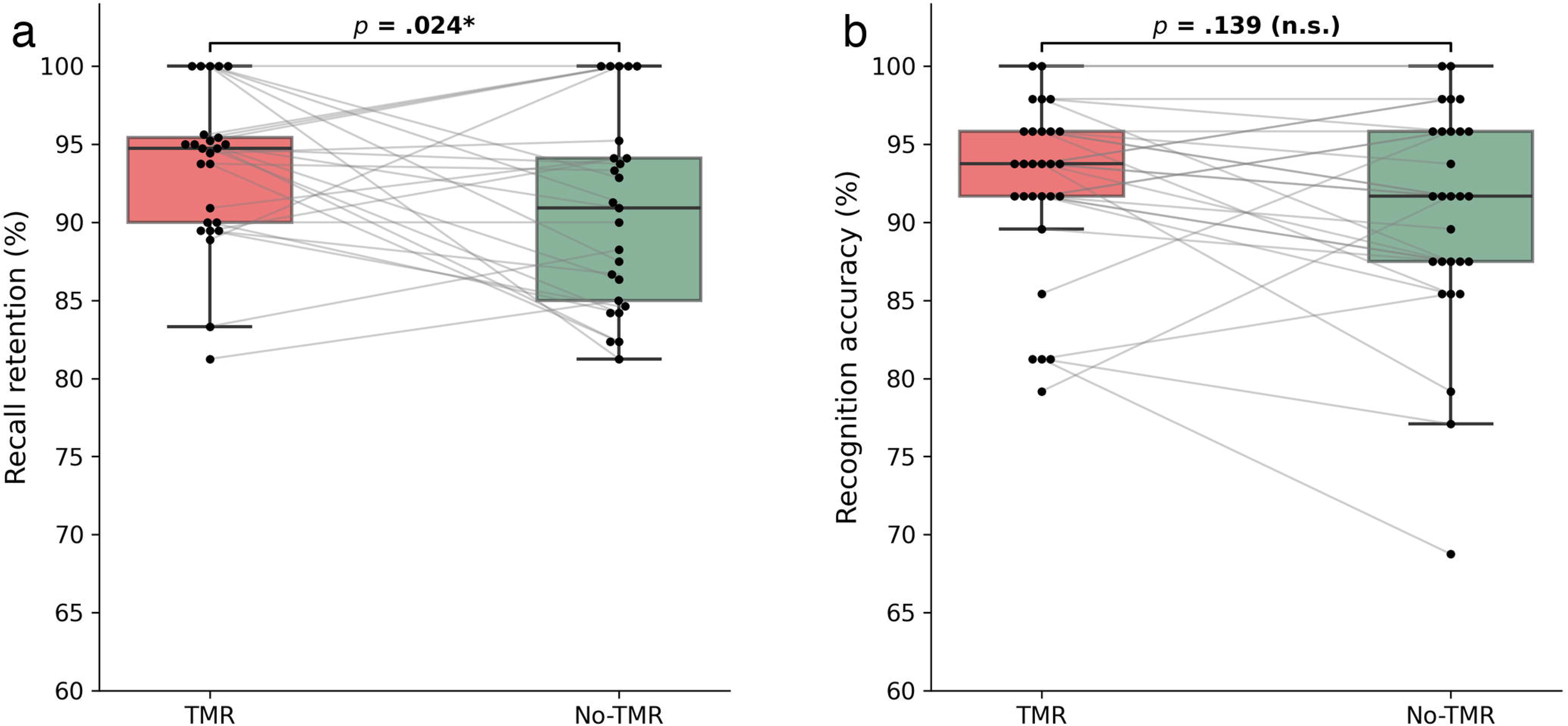
Post-sleep memory performance. Comparison of overnight memory performance between TMR (red) and No-TMR (green) conditions. (a) Recall retention rate (percentage of item locations correctly recalled). (b) Recognition accuracy [(hits + correct rejections)/48 × 100]. Boxes show the median and interquartile range; whiskers extend to the most extreme values within 1.5 × IQR. Individual data points represent single participants; paired lines connect individual participant scores across conditions. Effect sizes are Cohen’s *d* = 0.45 for recall and *d* = 0.29 for recognition. \**p* < .05; n.s. = not significant. **Alt text**. Two-panel boxplot comparison of post-sleep memory performance between TMR and No-TMR conditions across 28 participants. Panel a shows recall retention rate, with TMR slightly higher than No-TMR and the difference marked statistically significant. Panel b shows recognition accuracy, with closely similar distributions across the two conditions and the difference marked non-significant. Each panel displays box-and-whisker plots overlaid with individual participant data points connected by paired lines.

### EEG results

#### Sleep Architecture and Spindle Characteristics

On average, participants slept a total of 415.96 minutes (*SEM* = 6.94), with 215.39 minutes (*SEM* = 5.57) spent in NREM2 and 89.32 minutes (*SEM* = 5.44) in NREM3 sleep (see Table 2a for detailed sleep architecture). Offline spindle detection identified an average of 1658.43 ground-truth spindles per participant (*SEM* = 78.71). The Portiloop device delivered an average of 1107.21 stimulations across the night (*SEM* = 82.77), of which 817.18 (*SEM* = 56.42) were confirmed as true positive stimulations coinciding with validated spindles. Sleep spindle variables, including temporal organization, are summarized in Table 2b-d. Of the stimulated ground-truth spindles, 72.39% (*SEM* = 1.52) occurred within spindle trains and 27.61% (*SEM* = 1.52) occurred as isolated events. On average, 58.06% (*SEM* = 1.26) of all spindles within a given train were stimulated, indicating that closed-loop stimulation sampled the majority of train-embedded spindles. These spindle variables provide the basis for the correlational analyses reported below, which test whether the temporal context of stimulation differentially predicts memory outcomes.

#### Not all spindles are alike: Spindle temporal organization is selectively associated with distinct memory outcomes

To examine whether the temporal context of spindle-locked stimulation was associated with distinct memory outcomes, Spearman correlations were computed between spindle proportion variables and the TMR effect (TMR minus No-TMR) for both recall and recognition. To provide a complete test of specificity, all four combinations of spindle type (train, isolated) and memory outcomes (recall, recognition) are reported, along with Fisher *r*-to-*z* comparisons between spindle types for each outcome.

The proportion of spindles stimulated within a train was significantly and positively correlated with the TMR effect on recall, ρ (26) = .531, 95% CI [.20, .76], *p* = .004 (Pearson *r* = .452, 95% CI [.09, .71], *p* = .016), but not with the TMR effect on recognition, ρ (26) = .147, 95% CI [-.24, .49], *p* = .455 (Figure 5a). Conversely, the proportion of isolated spindles stimulated was significantly correlated with the TMR effect on recognition, ρ (26) = .563, 95% CI [.24, .77], *p* = .002 (Pearson *r* = .421, 95% CI [.06, .69], *p* = .026), but not with the TMR effect on recall, ρ (26) = .063, 95% CI [-.32, .43], *p* = .749 (Figure 5b). This pattern suggests a selective association between spindle temporal organization and memory outcome: stimulation during train-embedded spindles was associated with recall-based retention, whereas stimulation during isolated spindles was associated with recognition accuracy.

**Figure 5.**
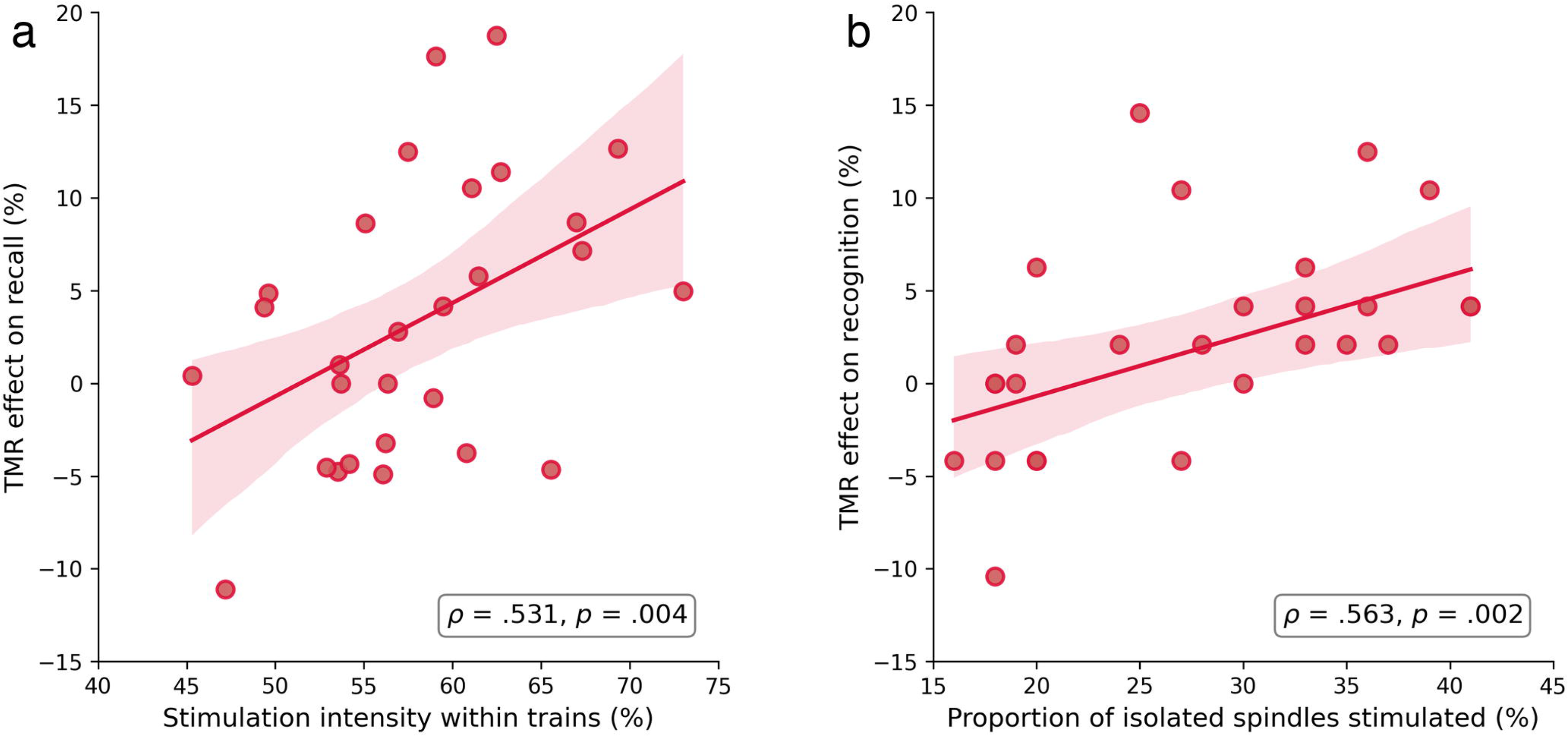
Spindle temporal organization and memory consolidation. (a) Correlation between stimulation intensity within trains (the average percentage of ground-truth spindles within each train that received stimulation, averaged across trains) and the TMR effect (TMR minus No-TMR) on recall. (b) Correlation between the proportion of isolated spindles stimulated and the TMR effect on recognition. Scatter plots show individual participant data with regression lines and 95% confidence intervals. Spearman correlation coefficients (ρ) and *p*-values are reported. **Alt text**. Two-panel scatter plot showing correlations between spindle stimulation patterns and the TMR memory effect across 28 participants. Panel a plots stimulation intensity within spindle trains on the x-axis against the TMR effect on recall on the y-axis, showing a positive correlation marked statistically significant. Panel b plots the proportion of isolated spindles stimulated against the TMR effect on recognition, showing a positive correlation also marked statistically significant. Each panel displays individual data points with a regression line and shaded 95 percent confidence interval.

Fisher *r*-to-*z* tests comparing the Pearson correlations between spindle types were not significant for recall (*z* = 1.693, *p* = .090) nor recognition (*z* = -1.296, *p* = .195). The observed pattern is therefore described as a selective association rather than a statistically confirmed double dissociation.

## Discussion

The present study examined whether sleep-dependent declarative memory consolidation can be modulated by TMR time-locked to sleep spindles using a closed-loop approach, and whether the temporal organization of the targeted spindles – grouped in trains versus isolated events – is associated with distinct memory outcomes. Behaviorally, post-sleep recall accuracy was significantly higher in the TMR than the No-TMR condition, whereas recognition accuracy and overnight change in movement time did not differ significantly between conditions. At the electrophysiological level, the proportion of spindles stimulated within a train was selectively associated with the TMR effect on recall, whereas the proportion of stimulated isolated spindles was selectively associated with the TMR effect on recognition. Together, these findings suggest that not all spindles contribute equally to memory consolidation and that the temporal context in which stimulated spindles occur is selectively associated with distinct memory outcomes.

The validity of these effects is supported by the fact that encoding performance improved reliably across successive blocks, with no difference between conditions and no Block x Condition interaction. This pattern indicates that repeated exposure produced incremental gains in performance ^80^, and that the learning trajectory was comparable across item categories later assigned to the TMR and No-TMR conditions. This equivalent encoding strength across conditions reduces the likelihood that post-sleep differences reflect pre-existing encoding biases rather than sleep-dependent consolidation mechanisms. Moreover, post-sleep testing revealed that item locations in the TMR condition were recalled at a significantly higher rate than those in the No-TMR condition, though the two conditions did not differ significantly on recognition accuracy or movement time. These findings converge with prior TMR work demonstrating that re-presenting sensory cues associated with learned material during NREM sleep facilitates consolidation of declarative memory ^1,42–44,52^. Importantly, our results also extend these findings by showing that recall-based retention benefits arise when cues are delivered time-locked to sleep spindles, though this effect did not extend uniformly to all memory measures. That spindle-locked TMR produced a significant recall effect here is consistent with the view that temporal alignment of cues to specific oscillatory events may be critical for effective memory reactivation. This contrasts with some prior open-loop TMR studies targeting broader NREM windows that have not consistently replicated behavioral gains ^81^.

At the group level, the TMR effect was significant for recall but not for recognition accuracy or movement time. This pattern suggests that spindle-locked TMR preferentially enhanced recall-based retention rather than strengthening memory representations uniformly across retrieval processes. One possible explanation is that the majority of stimulations were delivered during train-embedded spindles (72.39%), which were selectively associated with recall performance. By contrast, only 27.61% of stimulated spindles were isolated events – the spindle context selectively associated with recognition. The asymmetric distribution of stimulations across spindle types may thus have favored recall-related consolidation processes at the group level, while limiting the statistical power to detect a group-level recognition effect.

Consistent with this interpretation, the individual-level correlation between isolated spindle stimulation and the TMR recognition effect was significant (ρ = .563, *p* = .002), suggesting that recognition-related benefits may emerge when a significant proportion of stimulations coincide with isolated spindle events. Nonetheless, the observation that spindle-locked TMR enhanced recall extends previous work demonstrating that spindles provide a consolidation-supportive temporal window ^1,9^, and is consistent with accumulating evidence that auditory processing is not fully suppressed during spindle events (see ^59,60^ for a different view). Rather, externally presented sounds can be processed during spindles and interact with ongoing thalamocortical dynamics, allowing sensory cues to influence reactivation-related processes ^38,61,82^.

If spindles provide a temporal window favorable to memory consolidation, their temporal organization may further shape reactivation efficacy. Our correlational analyses revealed that spindle temporal context was selectively associated with different memory outcomes: stimulation intensity within trains – the proportion of train-embedded spindles that received stimulation – was associated with the TMR effect on recall, whereas the proportion of stimulated isolated spindles was associated with the TMR effect on recognition. Because these spindle proportion measures were derived within participants across a single stimulation session, the observed associations reflect within-subject variation in the temporal context of stimulation. Furthermore, because train-embedded and isolated stimulation represent complementary proportions of the same stimulation set, these associations should be interpreted as reflecting relative differences in temporal context rather than fully independent effects of each spindle type. It should also be noted that stimulated spindles in trains outnumbered isolated stimulated spindles. The possibility that this numerical imbalance, rather than a genuine difference in spindle-type function, contributes to the observed correlational pattern cannot therefore be fully excluded. Nonetheless, the current findings extend the emerging evidence that spindle temporal organization is functionally meaningful for consolidation ^8,26,83^ by demonstrating that distinct spindle contexts are selectively associated with different memory outcomes.

Mechanistically, spindle trains may provide temporally extended windows for iterative hippocampo-cortical communication, increasing the probability that auditory cues trigger successive reinstatement cycles that reinforce newly encoded memory traces. Such repeated reinforcement may explain why greater stimulation intensity within trains was associated with better recall-based retention. In contrast, isolated spindles may support more transient reinstatement episodes, sufficient to facilitate familiarity-based recognition – a comparatively less resource-demanding process ^70,71^ – while being less effective for strengthening the robust representations that underlie recollection-based recall. To our knowledge, the current study thus provides the first evidence that TMR using auditory cues time-locked to sleep spindles enhances declarative memory retention, while further showing that variability in TMR efficacy is associated with differences in the temporal organization of targeted spindles. These findings align with prior evidence that spindle temporal structuring is functionally relevant for memory reactivation and integration ^26,57,83^, motivating further mechanistic examination of how spindle timing governs the efficacy of TMR.

Building on this link between spindle timing and behavioral outcomes, our approach advances TMR methodology by time-locking auditory cues to ongoing spindles detected in real-time and further characterizing their occurrence in trains versus isolated events. This design builds on prior closed-loop studies that synchronized auditory stimulation with the slow oscillation up-state ^56,58,84,85^ but rarely examined the specific contribution of spindles or their temporal organization. Whereas most reports of spindle-memory associations have been correlational ^11,57,86^, our real-time closed-loop protocol allowed a more direct test of the relationship between spindle-locked stimulation and memory outcomes by experimentally delivering cues contingent on detected spindle events. This approach provides a more refined window into how spindle activity relates to reactivation processes, moving beyond purely correlational accounts of spindle-dependent memory consolidation.

While the current study provides novel insights into how spindle temporal organization relates to memory consolidation, several limitations should be acknowledged. First, and most critically, the present design compared spindle-locked TMR to a no-stimulation control. Without a condition in which TMR cues are delivered at random times during NREM sleep or in an open-loop fashion, it is not possible to determine whether the observed recall enhancement is specific to spindle-locked timing or whether similar benefits would emerge from any auditory stimulation during NREM2/3 sleep. Further studies should include such a control condition to directly test the added value of spindle-locked TMR over standard open-loop TMR. Second, the current work was conducted in a single-night design with healthy young adults. Extending these findings to older adults, children, or clinical populations with altered spindle dynamics will be important for establishing their generalizability. Third, we did not directly assess the neural content mediating the reactivation process, nor did we examine whether evoked EEG responses to auditory cues differed between stimulations delivered during spindle trains versus isolated spindles. Future work combining spindle-targeted TMR with EEG-fMRI, multivariate decoding, or time frequency analyses of cue-evoked responses could clarify the neural mechanisms underlying the observed behavioral dissociation. Fourth, the present study did not examine whether stimulated spindles were nested within slow oscillation up-states, a factor known to modulate consolidation efficacy as well ^56,87^. Given that spindle-SO coupling is thought to optimize the temporal alignment of memory reactivation with cortical excitability windows, future work should assess whether the memory benefits of spindle-train stimulation are further enhanced when cues coincide with SO-coupled spindle events. Fifth, our stimulation protocol used a minimum 3.5 s inter-stimulation interval, following prior work on the spindle refractory period ^57^ and pilot data from our own laboratory ^88^. While this conservative spacing minimized interference between consecutive cues, it also resulted in some train-embedded spindles not receiving stimulation. Future implementations could incorporate adaptive algorithms that dynamically adjust stimulation timing based on ongoing spindle density ^56,89^, thereby optimizing stimulation coverage within trains. Finally, the correlational nature of the spindle train and isolated spindle associations precludes causal conclusions about the role of spindle temporal organization per se; future experimental designs that selectively target trains versus isolated spindles could provide a more direct test of this relationship.

In conclusion, the present study demonstrates that TMR using auditory cues delivered in real-time during sleep spindles enhances declarative memory retention in the TMR relative to the No-TMR condition. Furthermore, the temporal context of stimulation was selectively associated with distinct memory outcomes, with train-embedded stimulation linked to recall and isolated stimulation linked to recognition. These findings highlight spindle temporal organization as a functionally relevant dimension of sleep-dependent consolidation and suggest that not all spindles contribute equally to memory. By targeting spindles in real time with closed-loop TMR, this work advances our understanding of how spindle dynamics support memory consolidation and provides a foundation for developing more precise, spindle-informed interventions.

Extending this approach to clinical populations with altered spindle dynamics may further reveal how disruptions in sleep oscillatory architecture contribute to memory impairment, highlighting the potential of spindle-based measures as clinically relevant biomarkers of brain health.

## Author’s Contributions

*Vaishali Mutreja*: Writing – original draft, Funding acquisition, conceptualization, data collection, methodology, formal analysis, figure generation, data curation; *Prakriti Gupta*: conceptualization, data collection, writing – review and editing; *Ovidiu Lungu*: conceptualization, methodology, data collection, data analysis, writing – review and editing; *Latifa Lazzouni*: conceptualization, methodology, data collection, data analysis, writing – review and editing; *Ella Gabitov*: conceptualization, methodology, data collection, writing – review and editing; *Habib Benali*: writing – review and editing; *Hugo R. Jourde*: methodology, writing – review and editing; *Giovanni Beltrame*: methodology, writing – review and editing; *Emily B.J. Coffey*: methodology, writing – review and editing; *Jean-Marc Lina*: writing – review and editing; *Genevieve Albouy*: writing – review and editing; *Bradley R. King*: writing – review and editing; *Arnaud Boutin*: writing – review and editing; *Julie Carrier*: writing – review and editing; *Julien Doyon*: funding acquisition, conceptualization, methodology, writing – review and editing, supervision.

## Funding

This work was supported by a Canadian Institutes of Health Research (CIHR) grant (#254572) to J.D.; the Jeanne Timmins Costello Studentship to V.M.; and a Healthy Brains, Healthy Lives (HBHL) doctoral fellowship to V.M.

## Acknowledgements

We are grateful to Hannah James for her assistance with data collection and to all the participants for their time and dedication to this study. We also thank the Portiloop engineering and research team for their technical support and guidance during device implementation and data acquisition.

## Use of AI Tools

During the preparation of this work, the authors used ChatGPT (OpenAI) to improve the clarity and readability of the manuscript. After using this tool, the authors reviewed and edited the content as needed and take full responsibility for the content of the publication.

## Disclosure Statement

Financial Disclosure: none. Non-financial Disclosure: none.

The manuscript will be deposited as a preprint to bioRxiv following submission. It has not been previously published and is not under consideration elsewhere.

## Data Availability Statement

The data that support the findings of this study are not publicly available due to participant confidentiality requirements set by the Institutional Review Board at McGill University (22-02-45, A03-B21-22A). Reasonable requests to access the data will be considered on a case-by-case basis and should be made to the corresponding author. Ethical approval for data sharing agreements is required to share data, in order to protect participant confidentiality.

## Notes

### Competing Interest Statement

The authors have declared no competing interest.

