## Supplementary file for "Sleep Spindle-Locked Targeted Memory Reactivation Enhances Declarative Memory Consolidation"

Supplementary Methods and Results

**Authors**

Vaishali Mutreja[1,2], Prakriti Gupta[1,2], Ovidiu Lungu[2,3], Latifa Lazzouni[4], Ella Gabitov[2,4], Habib Benali[5], Hugo R Jourde[6], Giovanni Beltrame[7], Emily B.J. Coffey[6], Jean-Marc Lina[8], Genevieve Albouy[9], Bradley R. King[9], Arnaud Boutin[10], Julie Carrier[11]& Julien Doyon[2,4]

**Affiliation**

[1]Integrated Program in Neuroscience, McGill University, Montreal, Canada
[2]McConnell Brain Imaging Centre, McGill University, Montreal, Canada
[3]Department of Psychiatry and Addictology, University of Montreal, Montreal, Canada
[4]Department of Neurology and Neurosurgery, McGill University, Montreal, Canada
[5]Electrical and Computer Engineering, Concordia University, Montreal, Canada
[6]Department of Psychology, Concordia University, Montreal, Canada
[7]Department of Computer and Software Engineering, Polytechnique Montreal, Montreal, Canada
[8]Department of Electrical Engineering, École de technologies supérieures (ETS), Montreal, Canada
[9]Department of Health and Kinesiology, University of Utah, Utah, United States
[10]Université Paris-Saclay, Inria, CIAMS, Gif-sur-Yvette, France
[11]Department of Psychology, University of Montreal, Montreal, Canada

To whom correspondence may be addressed

**List of Supplementary Materials**

**Supplementary Tables**

Table S1. Performance of the Portiloop spindle detection classifier across four probability thresholds.

Table S2. Participants' characteristics for the pilot stimulation timing study.

Table S3. Sleep architecture during habituation night.

Table S4. Block-by-block category means for encoding performance by stimulus category.

**Supplementary Figures**

Figure S1. Portiloop device classifier threshold validation.

Figure S2. Overnight closed-loop device performance.

Figure S3. Timing of the stimulation: individual overnight changes in recall accuracy and movement time.

### 1. Supplementary Methods

#### 1.1 Screening questionnaires

Eligibility criteria included right-handedness assessed using the Edinburgh Handedness Inventory [1], self-reported normal hearing, and the absence of factors known to influence sleep or cognition. Specifically, participants were excluded if they were smokers, used recreational drugs, had a history of medical, neurological, psychological, or psychiatric conditions (including depression and anxiety; participants scoring ≥ 8 on the Beck Depression Inventory [BDI; Beck, Rial and Rickels [2]] or the Beck Anxiety Inventory [BAI; Beck, Epstein, Brown and Steer [3]] were excluded), reported sleep disturbances (Pittsburgh Sleep Quality Index global score ≥ 5; Buysse, Reynolds, Monk, Berman and Kupfer [4]), or were taking any psychoactive or sleep-affecting medications.

To minimize the impact of circadian and sleep-related variability, participants were instructed to maintain a regular sleep-wake schedule (bedtime: 22:00-23:30; wake time: 07:00-08:30) for at least three days before the experiment. They were also asked to abstain from alcohol and caffeine for 24 hours prior to the experimental session.

#### 1.2 Auditory stimuli and calibration

Two auditory stimuli were used throughout the experiment: (1) complex harmonic tone and (2) a spoken vowel ("A"). The harmonic tone has a fundamental frequency of 523 Hz (musical tone C5) and consisted of 12 harmonics with linearly decreasing amplitude. The waveform was synthetically generated to ensure spectral stability and uniformity across presentations. The vowel stimulus consisted of a naturally produced phoneme ("A"). Both stimuli were 100 ms in duration and sampled at 44.1 kHz. A 10-ms linear ramp was applied at stimulus onset and offset to prevent spectral splatter and audible transients. Stimuli were normalized prior to calibration to ensure consistent relative intensity across participants.

Auditory stimuli presented during the OSL task inside the MRI scanner were delivered via MR-compatible earphones, whereas stimuli presented during sleep were delivered through Etymotic ER 3C insert earphones with foam tips. Individual auditory detection thresholds for each stimulus were determined for each participant using a one-down/one-up staircase procedure while participants lay supine in the MRI scanner. Based on these thresholds, sound intensity was adjusted to a level that was clearly audible during task performance while remaining comfortable and non-intrusive.

Prior to the overnight sleep session, sound levels were adjusted to ensure audibility without inducing arousal or awakening. Sound pressure levels were calibrated using a sound level meter to confirm consistent stimulation device output delivery across sound cues. Mean volume intensities were 34.62 dB (SD = 24.45) for the harmonic tone and 45.00 dB (SD = 23.15) for the vowel stimulus.

Category-sound pairings were counterbalanced across participants to avoid potential confounds related to sound-category associations established during pre-sleep learning. The same calibrated stimulus parameters were used across encoding, feedback, and sleep stimulation to maintain consistency in auditory exposure.

#### 1.3 Analyses

##### 1.3.1 EEG data Analysis

**EEG preprocessing**: The preprocessing pipeline was implemented using EEGLAB [5], and custom MATLAB scripts and was applied on the raw EEG signals collected from the Portiloop device. EEG preprocessing steps included mainly: application of a broadband filter (0.3 – 30 Hz) and artifact detection.

Data import and setup: The raw EEG signals were imported from the Portiloop system, containing continuous EEG data. The data was loaded into EEGLAB for further processing.

Filtering: A bandpass filter (0.3 – 30 Hz) was applied to the data to remove unwanted noise outside the frequency range of interest. This filtering was performed using an FIR filter with a filter order of 100, optimal for the data. The filtering process helped isolate the frequency bands of interest, enhancing the signal-to-noise ratio for subsequent analyses.

Artifact detection: A continuous artifact rejection approach was employed to eliminate noisy segments in the data. This process marked intervals with significant artifacts, such as high-amplitude spikes, and excluded them from the final dataset. The artifact rejection was conducted based on a peak-to-peak threshold, set at 400 µV for the channels of interest, and utilizing a moving window approach (window size: 200 ms, step size: 100 ms) to identify and reject segments with excessive artifact contamination. The basicrap function (BASIC artifact rejection by peak-to-peak; ERPLAB) was used to identify and mark the bad segments rather than cutting them from the dataset. The marked artifacts were saved as specific events in the EEG data structure as bad intervals.

After preprocessing, the cleaned/marked EEG data was saved in EEGLAB with events associated with bad intervals marked for later review.

**Spindle clustering**: After performing the spindle detection procedure, sleep spindles were classified as occurring in trains or as isolated events based solely on their temporal organization within the EEG signal, independent of auditory stimulation, following the framework proposed by Boutin and Doyon [6]. Spindle trains were defined as sequences of spindles occurring within 6s of one another, whereas isolated spindles were defined as spindles separated by more than 6s from both preceding and subsequent spindles, based on offline spindle detection.

Real-time stimulation events were then mapped onto offline ground truth spindle classification. Because stimulation delivery was constrained by a minimum inter-stimulation interval to account for the spindle refractory period, not all detected spindles within a train were necessarily stimulated. Each stimulated spindle, based on the offline detection, was therefore classified as either occurring within a spindle train or occurring in isolation. Thus, a stimulated spindle was considered part of a stimulated spindle train if it occurred within a train defined by offline detection, even if adjacent spindles in the same train were not stimulated. Conversely, a stimulated spindle was classified as an isolated stimulated spindle if it occurred as an isolated event according to the offline spindle classification.

This approach ensured that spindle clustering reflected intrinsic spindle dynamics rather than stimulation timing and allowed us to examine how TMR delivered during spindles embedded in trains versus isolated spindles differentially influenced memory consolidation.

#### 2. Supplementary Results

##### 2.1 Closed-loop device threshold validation

**Validation of classifier probability threshold (pilot nap study)**

To determine an appropriate classifier probability threshold for real-time spindle detection, we conducted a pilot validation study using daytime nap recordings. This validation was performed to optimize the performance of the closed-loop stimulation device and was independent of the specific memory task used in the main experiment.

The classifier probability threshold defined the confidence level required by the Portiloop device [7] to label an event as a sleep spindle. Lower thresholds favor sensitivity by increasing the likelihood of detecting true spindles, whereas higher thresholds prioritize specificity at the cost of missing true spindle events. Selecting an optimal threshold therefore requires balancing detection specificity against false-positive stimulations.

Twelve nap recordings were initially collected at the 'Institut universitaire de gériatrie de Montréal (IUGM)' sleep laboratory from healthy young adults (18-30 years). Participants were screened using the same criteria as in the main study (see Methods), including absence of neurological or psychiatric disorders and abstinence from caffeine prior to recording. EEG data were visually scored according to AASM criteria by a certified sleep technologist. Offline spindle detection was performed at electrode Cz using the algorithm described by Wamsley, Tucker, Shinn, Ono, McKinley, Ely, Goff, Stickgold and Manoach [8], which served as a ground-truth reference. EEG preprocessing included band-pass filtering, rejection of noisy segments, and restriction of analyses to NREM2 and NREM3 sleep. Four recordings were excluded due to poor signal quality or insufficient sleep duration, leaving eight datasets for analysis.

Using the Portiloop demonstration/simulation framework, spindle detections were simulated at three classifier probability thresholds (0.71, 0.75, and 0.84) and compared against detections obtained during real-time acquisition at a threshold of 0.82. For each dataset, true positives, false positives, and false negatives were identified relative to the offline ground truth, and precision, sensitivity, and F1-scores were computed.

Across thresholds, mean precision ranged from 0.73 to 0.80, sensitivity from 0.48 to 0.67, and F1-score from 0.59 to 0.70 (Table S1; Figure S1). Thresholds of 0.71 and 0.75 showed comparable overall performance across all metrics. Given this equivalence, a threshold of 0.75 was selected for subsequent overnight experiments to provide a consistent operating point for closed-loop stimulation.

This classifier threshold was subsequently used in the main declarative memory experiment described in the present manuscript.

**Table S1**. *Performance of the Portiloop spindle detection classifier*

| Threshold | Precision (M±SD) | Sensitivity (M±SD) | F1-score (M±SD) |
|---|---|---|---|
| 0.71 | 0.74 ± 0.09 | 0.67 ± 0.10 | 0.70 ± 0.05 |
| 0.75 | 0.75 ± 0.08 | 0.64 ± 0.12 | 0.68 ± 0.07 |
| 0.84 | 0.81 ± 0.09 | 0.53 ± 0.12 | 0.64 ± 0.09 |
| 0.82 | 0.80 ± 0.08 | 0.48 ± 0.09 | 0.60 ± 0.08 |

**Table S1**. Performance of the Portiloop spindle detection classifier across four probability thresholds (0.71, 0.75, 0.84, 0.82). Values represent group means (M) ± standard deviation (SD) for precision, sensitivity, and F1-score (N = 8 nap recordings)

**Figure S1**.

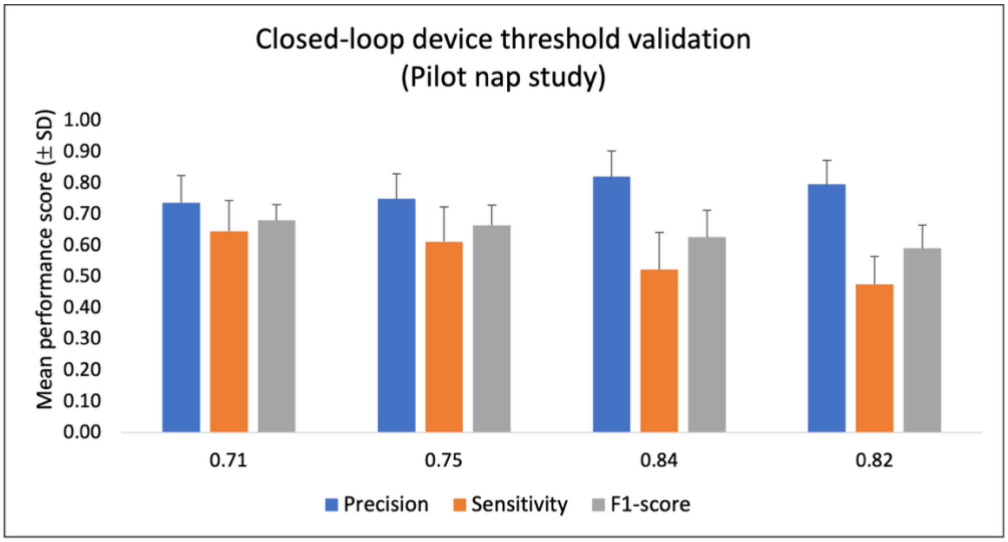

**Figure S1. Portiloop device classifier threshold validation**. Mean precision, sensitivity, and F1-scores (±SD) obtained during a pilot nap study (N = 8) across four classifier probability thresholds (0.71, 0.75, 0.84, 0.82). Spindles were detected offline using the Wamsley algorithm to establish ground-truth, and Portiloop detections were compared to compute true positives, and false negatives. Both 0.71 and 0.75 thresholds showed comparable performance; 0.75 was selected for subsequent overnight experiments.

#### 2.2 Overnight closed-loop device performance

To assess the reliability of the closed-loop stimulation system during full-night recordings, real-time spindle detections generated by the Portiloop were compared with offline visual spindle annotations obtained from the Cz electrode, selected as a representative central scalp site. Device performance was evaluated using three complementary indices: specificity (precision), sensitivity and F1-score.

Precision (positive predictive value) was defined as the proportion of real-time detections that were confirmed visually, indexing the accuracy with which auditory cues were delivered during true spindle events. Sensitivity (true positive rate) quantified the proportion of visually identified spindles that were successfully detected online. To capture the trade-off between these two measures, an F1-score was computed as the harmonic mean of precision and sensitivity.

Across participants, the Portiloop achieved a mean precision of 0.74±0.07, indicating that the majority of detected events corresponded to bona fide spindles. Sensitivity was lower at 0.59±0.15, reflecting that a subset of visually scored spindles were not detected in real time. The resulting F1-score averaged 0.65±0.09, consistent with a conservative detection strategy favoring specificity over maximal spindle capture.

Overall, these results demonstrate stable and reliable real-time spindle detection across overnight recordings. This performance profile ensured that most auditory cues were delivered during genuine spindle activity, supporting the validity of the closed-loop stimulation protocol used in the declarative memory experiment (Figure S2).

**Figure S2**.

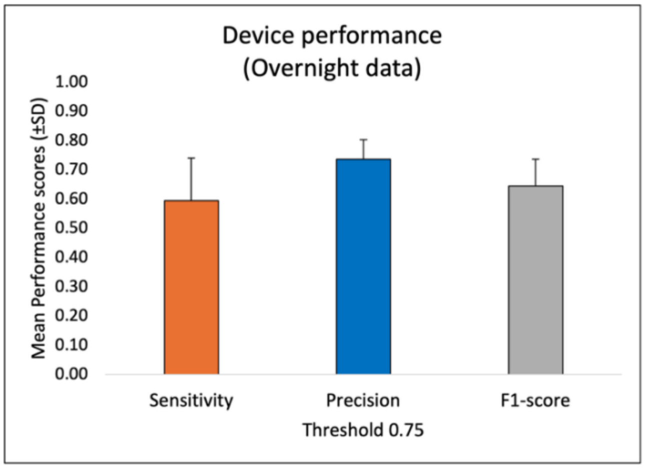

**Figure S2**. **Overnight closed-loop device performance**. Portiloop device performance for overnight recordings of the present study at the finalized threshold of 0.75 (N = 28). Mean precision, sensitivity, and F1-scores (±SD) are shown. The device demonstrated reliable spindle detection across full-night EEG recordings.

##### 2.3 Targeted Memory Stimulation Protocol: Timing of the stimulation

A 2018 study by Antony and colleagues [9] demonstrated that spindles are followed by a refractory period of approximately 0.5-2.5 s during which new spindles are less likely to occur. Using real-time spindle tracking, they showed that TMR cues presented after this refractory period improved memory performance more effectively than cues delivered within it. As closed-loop auditory stimulation time-locked to sleep spindles remains a novel approach, we conducted a pilot study to compare the behavioral effects of delivering stimulation during spindles versus outside the spindle refractory period.

The study design was similar to the main experiment described in the main manuscript, with the difference that instead of having a within-subject approach, we used a mixed between-within design. Specifically, we collected data from twelve participants (mean age: 26 years, range: 21 – 29) divided into two groups: six in the 'during spindle' group and six in the 'outside refractory period' group (see Table S2) and we manipulated the TMR vs. No-TMR condition within each. In the 'during spindle' group, TMR auditory cues were delivered as soon as a spindle was detected, followed by a minimum inter-stimulation interval of 3.5 s, based on Antony, Piloto, Wang, Pacheco, Norman and Paller [9]. In the 'outside refractory period' group,

TMR stimulation was triggered 3.5 s after spindle onset to ensure cues were delivered beyond the refractory period.

**Table S2**. *Participant's characteristics*

| Group | During spindle | Outside refractory |
|---|---|---|
| Total Participants | 6 | 6 |
| Females | 5 | 2 |
| Males | 1 | 4 |
| Age-range | 22-29 | 21-28 |
| $m_{age}$ (SD) | 25.00 (2.79) | 23.83 (2.64) |
| $m_{age}$ (SD) Female | 26.00 (2.59) | 24.5 (4.95) |
| $m_{age}$ (SD) Male | 22.00 | 23.50 (1.73) |

*Notes*. Sex was unequally distributed across pilot groups due to availability constraints during the recruitment window. Given the descriptive nature of this pilot, no sex-balanced inferential analyses were conducted. Values are in years. $m_{age}$ = mean age of the participant; SD = Standard deviation.

Participants arrived at the sleep laboratory around 21:00h, completed the same object-spatial location task as that described in the main manuscript, and then slept overnight while EEG recordings were collected and auditory stimulations were delivered using the Portiloop closed-loop system (described in detail in the main study). Auditory stimuli were delivered through earphones during NREM sleep (~ 7-7.5 h total sleep). As in the main study, only one of the two category-specific sounds was reactivated during sleep; however, the timing was different across the two groups, with auditory stimulation delivered either during spindles or outside the refractory period, depending on group assignment.

As in the main study, behavioral performance was quantified as the overnight memory change (post-sleep minus pre-sleep) in recall accuracy (percentage of correctly recalled items) and standardized movement time for correct responses. Given the exploratory nature of this pilot study and the small sample size, no inferential statistical analyses were conducted. Instead, results are reported descriptively at the individual level.

In the ‘during spindle’ group, 5 out of 6 participants showed increase in recall accuracy for TMR items relative to No-TMR items, whereas 1 participant showed no change (Figure S3A). Similarly, 4 out of 6 participants exhibited faster standardized movement times for TMR items compared to No-TMR items, with the remaining participants showing minimal or no improvement (Figure S3B). Mean overnight accuracy change for TMR items in this group was positive, whereas No-TMR items showed little to no improvement.

In contrast, in the ‘outside refractory period’ group, no consistent benefits of TMR were observed. Accuracy changes for TMR items were either unchanged or decreased relative to No-TMR items in 4 out of 6 participants (Figure S3C). Standardized movement time showed substantial inter-individual variability, with no systematic improvement for TMR items at the group level (Figure S3D).

Overall, individual-level behavioral trajectories indicated a clear qualitative difference between stimulation delivered ‘during spindles’ and stimulation delivered ‘outside the refractory period’, with the former showing more consistent overnight improvements in both accuracy and movement efficiency. Based on these descriptive pilot results, the main experiment adopted a ‘during spindle’ stimulation protocol with a minimum inter-stimulation interval of 3.5 s.

**Figure S3**.

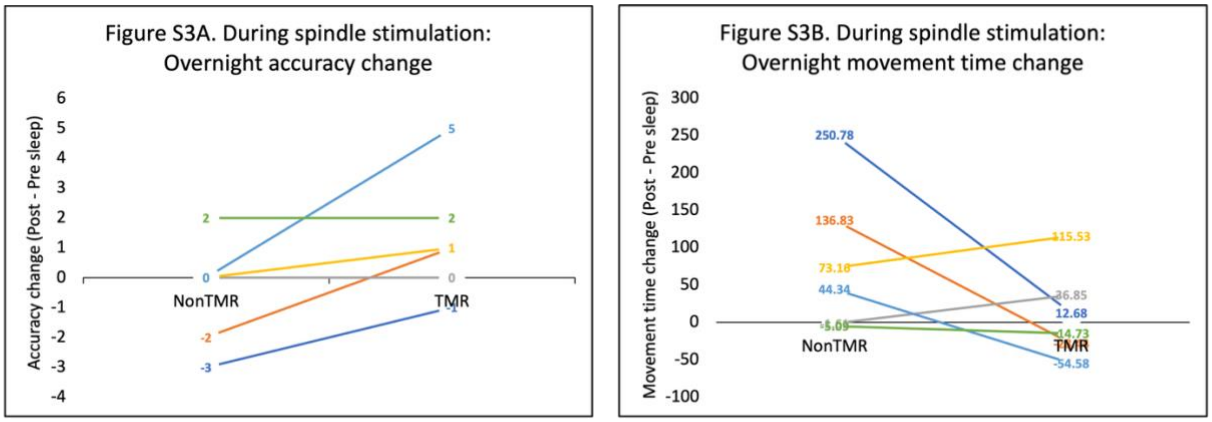

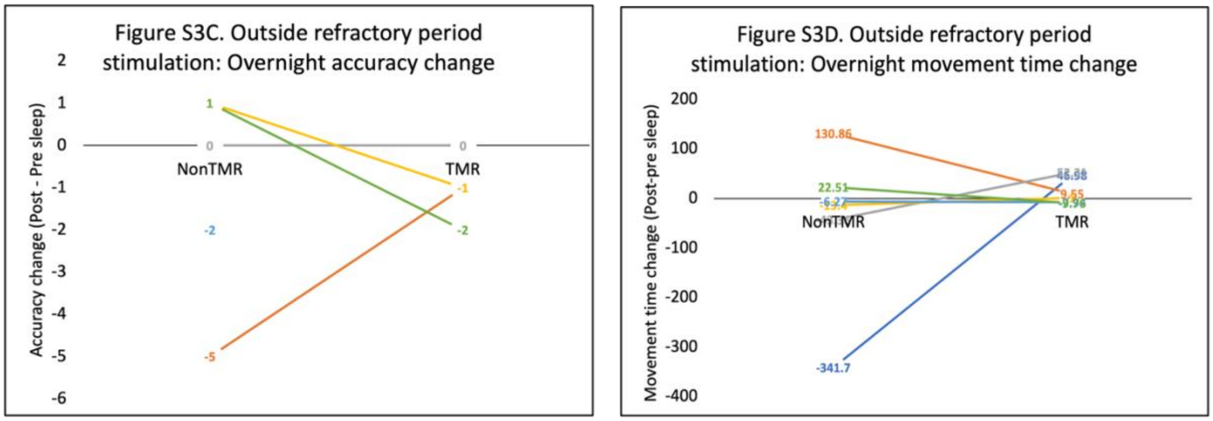

**Figure S3. Timing of the stimulation**. Individual overnight changes in recall accuracy (A,C) and standardized movement time (B, D) for TMR and No-TMR items in the 'during spindle' (A-B) and 'outside refractory period' (C-D) groups. Each line represents one participant. Values reflect post-sleep minus pre-sleep performance. Results are shown descriptively due to the pilot nature of the study.

##### 2.4 Habituation night sleep architecture

**Table S3**. *Sleep architecture during habituation night*

| Sleep stage | Mean ± SD | MIN | MAX |
|---|---|---|---|
| Total sleep time | 413.40 ± 49.09 | 230.50 | 470.00 |
| NREM1 | 20.88 ± 13.48 | 4.50 | 67.50 |
| NREM2 | 225.30 ± 44.65 | 99.00 | 297.50 |
| NREM3 | 82.70 ± 25.86 | 35.50 | 137.00 |
| REM | 84.52 ± 26.12 | 25.50 | 124.00 |

*Notes*. Descriptive statistics (mean ± SD, minimum, and maximum) for total sleep time and time spent in each sleep stage during habituation night, based on N = 25 participants (3 of 28 excluded due to processing issues). Sleep stages were scored according to standard criteria and are reported in minutes. NREM = non-rapid eye movement; REM = rapid eye movement.

The EEG recordings from the habituation night were reviewed by a certified sleep technologist to screen for any sleep abnormalities or disorders. Participants who passed this screening were invited for the experimental visit. Sleep architecture is summarized in Table S3. On average, participants slept for 413.40 minutes (SD = 49.09; N = 25). Data from three participants were excluded from the sleep architecture analysis due to processing issues; however, no abnormal sleep signs were reported by the sleep expert.

##### 2.5 Encoding performance by stimulus category

To assess whether encoding performance differed between stimulus categories (animals vs. clothing), a linear mixed-effects model was conducted with Block (1-4) and Category (animals, clothing) as within-subject factors. Results revealed a significant main effect of Block, confirming that recall accuracy improved across successive encoding blocks. There was no significant main effect of Category, indicating that encoding performance did not differ between animals and clothing items. The Block x Category interaction was also not significant. A paired-samples *t*-test comparing last-block performance confirmed no significant difference between animals ($M = 19.11$, $SEM = 0.33$) and clothing ($M = 18.75$, $SEM = 0.44$), $t(27) = 0.752$, $p = .458$. These results indicate that any post-sleep differences between TMR and No-TMR conditions are unlikely to reflect category-specific encoding advantages.

**Table S4**: *Block-by-block category means*

| **Block** | **Animals M (SEM)** | ***N*** | **Clothing M (SEM)** | **n** |
|---|---|---|---|---|
| 1 | 7.43 (0.87) | 28 | 6.00 (0.91) | 28 |
| 2 | 14.04 (0.89) | 27 | 13.15 (0.95) | 27 |
| 3 | 16.71 (1.02) | 21 | 15.38 (1.28) | 21 |
| 4 | 19.00 (0.65) | 9 | 17.22 (0.40) | 9 |
| Last block | 19.11 (0.33) | 28 | 18.75 (0.44) | |

*Notes*. Mean (SEM) recall accuracy across encoding blocks for animals vs. clothing categories. *N* = number of participants completing each block. Decreasing *N* across blocks reflects the criterion-based encoding procedure: participants who reached the ≥ 15/24 per-category criterion stopped before completing all four blocks. The Last block" row reports performance during each participant's final completed block (N = 28).
